# Chromatin at X-linked repeats that guide dosage compensation in *Drosophila melanogaster* is modulated by the siRNA pathway

**DOI:** 10.1101/330423

**Authors:** Nikita Deshpande, Victoria H. Meller

**Affiliations:** Department of Biological Sciences, Wayne State University, Detroit Michigan 48202

**Author notes:** Communicating author: 5047 Gullen Mall Wayne State University Detroit MI 48202.

**Keywords:** *Ago2*, dosage compensation, chromatin modification, satellite repeats, 1.688^X^ repeats, 359 bp repeats, *roX1 roX2*, X chromosome recognition

## Abstract

Many heterogametic organisms adjust sex chromosome expression to accommodate differences in gene dosage. This requires selective recruitment of regulatory factors to the modulated chromosome. How these factors are localized to a chromosome with requisite accuracy is poorly understood. *Drosophila melanogaster* males increase expression from their single X chromosome. Identification of this chromosome involves cooperation between different classes of X-identity elements. The Chromatin Entry Sites (CES) recruit a chromatin-modifying complex that spreads into nearby genes and increases expression. In addition, a family of satellite repeats that is enriched on the X chromosome, the 1.688^X^ repeats, promotes recruitment of the complex to nearby genes. The 1.688^X^ repeats and CES are dissimilar, and appear to operate through different mechanisms. Interestingly, the siRNA pathway and siRNA from a 1.688^X^ repeat also promote X recognition. We postulate that siRNA-dependent modification of 1.688^X^ chromatin contributes to recognition of nearby genes. In accord with this, we found enrichment of the siRNA effector Argonaute2 (Ago2) at some 1.688^X^ repeats.Mutations in several proteins that physically interact with Ago2, including the histone methyltransferase Su(var)3-9, enhance the lethality of males with defective X recognition. Su(var)3-9 deposits H3K9me2 on some 1.688^X^ repeats, and this mark is disrupted upon ectopic expression of 1.688^X^ siRNA. Furthermore, integration of 1.688^X^ DNA on an autosome induces local H3K9me2 deposition, but enhances expression of nearby genes in a siRNA-dependent manner. Our findings are consistent with a model in which siRNA-directed modification of 1.688^X^ chromatin contributes to recognition of the fly X chromosome by the MSL complex.

## Introduction

Males of many species carry one X chromosome and a gene-poor Y chromosome. Hemizygosity of the male X chromosome produces a potentially lethal imbalance in the ratio of X to autosomal gene products. This imbalance is corrected by a process known as dosage compensation, a specialized type of gene regulation that modulates expression of an entire chromosome. Different strategies to achieve dosage compensation have evolved independently. In *Drosophila melanogaster*, males increase X-linked gene expression by approximately two-fold (LUCCHESI *et al.* 2005).This involves the activity of the Male Specific Lethal (MSL) complex. The MSL complex is recruited to active genes on the X chromosome, where it modifies chromatin to increase expression (LUCCHESI AND KURODA 2015). The MSL complex contains five proteins, Male-Specific Lethal 1, -2, and -3 (MSL1, -2, -3), Maleless (MLE), and Males Absent on the First (MOF) (reviewed in (KOYA AND MELLER 2011)). Enhanced transcription by the MSL complex is associated with H4K16 acetylation by MOF (AKHTAR AND BECKER 2000; SMITH *et al.* 2000). H4K16 acetylation decompacts chromatin, and this may enhance transcriptional elongation of X-linked genes (SHOGREN-KNAAK *et al.* 2006; LARSCHAN *et al.* 2011).

The MSL complex also contains one of two non-coding *RNA on the X* (*roX1, -2*) transcripts (QUINN *et al.* 2014). While elimination of any one of the MSL proteins is lethal to males, *roX1* and *roX2* are redundant for compensation. Mutation of both *roX* genes leads to mislocalization of the MSL proteins to ectopic autosomal sites in male larvae (MELLER AND RATTNER 2002; DENG AND MELLER 2006). X-linked gene expression is reduced in these males, as is survival to adulthood. Both *roX* genes are located on the X chromosome, and both overlap Chromatin Entry Sites (CES), specialized sites with increased affinity for the MSL complex (KELLEY *et al.* 1999; ALEKSEYENKO *et al.* 2008; STRAUB *et al.* 2008).

Although much is known about the role of MSL complex in dosage compensation, how this complex selectively targets the X chromosome is poorly understood. Recognition and binding to X chromatin is believed to be a two-step process. Initial recruitment of the MSL complex to CES is followed by spreading into nearby transcribed genes (GELBART AND KURODA 2009). Contained within the CES are motifs called MSL Recognition Elements (MREs) (ALEKSEYENKO *et al.* 2008; STRAUB *et al.* 2008). MREs are 21 bp GA-rich motifs that bind Chromatin-Linked Adaptor for MSL Protein (CLAMP), a zinc finger protein that is essential for MSL recruitment (SORUCO *et al.* 2013).Spreading into nearby active genes is supported by interaction of MSL3 with the cotranscriptional H3K36me3 mark (KIND AND AKHTAR 2007; LARSCHAN *et al.* 2007; SURAL *et al.* 2008). These mechanisms describe local recruitment of the MSL complex, but fail to explain how the MSL complex specifically targets the X-chromosome. H3K36me3 is found on active genes throughout the genome, and MREs are only modestly enriched on the X chromosome. Furthermore, CLAMP binds MREs throughout the genome, but only recruits the MSL complex to X-linked CES (ALEKSEYENKO *et al.* 2008; SORUCO *et al.* 2013). We conclude that additional mechanisms must distinguish X and autosomal chromatin.

X-localization is disrupted in *roX1 roX2* males, making them a sensitized genetic background that can be used to identify additional factors contributing to X recognition.Using this strategy, our laboratory demonstrated a role for the siRNA pathway in recognition of the X-chromosome (MENON AND MELLER 2012; MENON *et al.* 2014). A likely source of siRNA is a family of repeats that is near exclusive to the X chromosome. These are the AT rich, 359 bp 1.688^X^ satellite repeats, a clade of which is found in short, tandem arrays throughout X euchromatin (HSIEH AND BRUTLAG 1979; WARING AND POLLACK 1987; DIBARTOLOMEIS *et al.* 1992; GALLACH 2014). Specific clusters are denoted by a superscript indicating cytological position. In support of the idea that 1.688^X^ repeats assist X recognition, ectopic production of siRNA from one repeat partially rescues *roX1 roX2* males (MENON *et al.* 2014). 1.688^X^ repeats are often close to or within genes, leading to the idea that they function as “tuning knobs” for gene regulation (KUHN *et al.* 2012). In accord with these ideas, autosomal insertions of 1.688^X^ DNA enable recruitment of functional dosage compensation to nearby autosomal genes (JOSHI AND MELLER 2017).

The 1.688^X^ repeats share no sequence identity with the CES, and appear to act in a genetically distinct manner (JOSHI AND MELLER 2017). The question of how 1.688^X^ DNA promotes compensation of nearby genes is thus of great interest. We pursued the idea that siRNA-directed modifications of chromatin at 1.688^X^ repeats link the repeats and the siRNA pathway to X recognition. Reduction of the siRNA-binding effector protein Argonaute 2 (Ago2) enhances the lethality of partial loss of function *roX1 roX2* mutations, and further reduces X-localization of MSL proteins (MENON AND MELLER 2012). We hypothesized that an Ago2-containing complex might localize to and modify 1.688^X^ chromatin in otherwise wild type flies. In accord with this idea, we find that Ago2 is enriched at 1.688^X^ repeats. Proteins interacting with Ago2 may also play a role in dosage compensation. To address this, we tested high confidence Ago2-binding proteins for genetic interactions with *roX1 roX2*, and found that mutations in several of these genes further reduced the survival of *roX1 roX2* males. Of particular interest is the H3K9 methyltransferase *Su(var)3-9*, which is responsible for enrichment of H3K9me2 at a subset of 1.688^X^ repeats. H3K9me2 enrichment is disrupted upon ectopic expression of 1.688^X^ siRNA. Chromatin flanking an autosomal insertion of 1.688^X^ DNA is enriched for H3K9me2, and enrichment is enhanced by ectopic expression of 1.688^X^ siRNA. In contrast to the repressive nature of H3K9me2, we find that expression of autosomal genes close to the 1.688^X^ transgene is increased in male larvae, and further elevated by ectopic production of 1.688^X^ siRNA. These findings support the idea that X recognition and transcriptional upregulation by dosage compensation are distinct processes, and propose that siRNA-dependent modification of chromatin in or near 1.688^X^ repeats contributes to X recognition in wild type flies. We propose that epigenetic modifications link the siRNA pathway, 1.688^X^ repeats on the X chromosome and X recognition.

## Materials and Methods

### Fly culture and Genetics

Mutations *Dcr1*^*Q1147X*^ (BDSC #32066), *Rm62*^*01086*^ (BDSC #11520), *Fmr1*^*Δ113m*^ (BDSC #67403), *Su(var)3-9*^*1*^ (BDSC #6209), S*u(var)3-9*^*2*^ (BDSC #6210), *smg*^*1*^ (BDSC #5930), *Taf11*^*1*^ (BDSC #65410), *Taf11*^*5*^ (BDSC #65409), *p53*^*5A-1-4*^ (BDSC #6815), *p53*^*11-1B-1*^ (BDSC #6816), *foxo*^*Δ94*^ (BDSC #42220), *PIG-S*^*e00272*^ (BDSC #17833), b*el*^*L4740*^ (BDSC #10222), *bel*^*6*^ (BDSC #4024), *barr*^*L305*^ (BDSC #4402), *SmD1*^*EY01516*^ (BDSC #15514), v*ig*^*C274*^ (BDSC #16323), *Ago1*^*k08121*^ (BDSC #10772), *aub*^*QC42*^ (BDSC #4968), *piwi*^*06843*^ (BDSC #12225), S*u(var)2-10*^*2*^ (BDSC #6235), *egg*^*MB00702*^ (BDSC #22876), *G9a*^*MB11975*^ (BDSC #29933), P{EPgy2}^*09821*^ (BDSC #16954), P{EPgy2}^*15840*^ (BDSC #21163), and *FLAG.HA.Ago2* (BDSC #33242)were obtained from the Bloomington Drosophila Stock Center. *Ago2*^*414*^ (Kyoto #109027) was obtained from the Kyoto Stock Center. *Su(var)3-7*^*14*^ was a gift from Dr. P. Spierer (SEUM *et al.* 2002). *ocm*^*166*^ was a gift from Dr. R. Kelley. *ΔDsRedΔupSET* (*upSET* in Figure 2) was a gift from Dr. M. Kuroda (MCELROY *et al.* 2017). To minimize genetic background effects all mutations were out crossed for five generations using a nearby marked *P*-element (unmarked mutations) or the laboratory reference *yw* strain (mutations marked with *w*^+^ or *y*^+^). Stocks were constructed with outcrossed, rebalanced chromosomes and a reference Y-chromosome (MENON AND MELLER 2009). All mutations were confirmed by phenotype or PCR. Mating schemes to determine the effect of Ago2-interactors on dosage compensation are presented in Figure S1. Each test scored about 1000 flies and was performed in triplicate. To express 1.688^3F^ siRNA in a *Su(var)3-9*^*-/-*^ mutant background, we generated [*hp1.688*^*3F*^] [*Sqh*-Gal4]*/In*(2LR)*Gla wg*^*Gla-1*^; *Su(var)3-9*^*1*^/ *™3TbSb* flies and selected non-*Tb* third instar males for ChIP. The [*Sqh*-Gal4] insertion was a gift of Dr.S. Todi. The [*hp1.688*^*3F*^] transgene contains part of the 1.688^3F^ repeat cluster coned in inverted orientation in pWIZ (SIKLEE AND CARTHEW 2003). Although siRNA accumulates to abnormally high levels in larvae expressing [*hp1.688*^*3F*^], the siRNAs produced appear similar to those isolated from wild type embryos (MENON *et al.* 2014)

### Tissue collection and chromatin preparation

Embryo collection and chromatin preparation was as previously described (KOYA AND MELLER 2015). Briefly, 0.5 g of 0 - 12 hr embryos were collected on molasses plates with yeast. Embryos were dechorionated for 2.5 min in bleach, crosslinked in 50 mM HEPES, 1 mM EDTA, 0.5 mM EGTA, 100 mM NACl, 1 % formaldehyde with heptane for 20 min. Crosslinking was quenched with 125 mM glycine, 0.01 % Triton X-100, 1 X PBS for 30 min. Embryos were washed with 10 mM HEPES, 200 mM NaCl, 1 mM EDTA, 0.5 mM EGTA and 0.01 % Triton X-100 and suspended in 2.5 ml of 10 mM HEPES, 1 mM EDTA, 0.5 mM EGTA, 0.1 % Na-deoxycholate and 0.02 % Na-azide for sonication. Sonication, performed on ice at 35 % amplitude, 30 sec on, 59 sec off for a total time 15 min using a Fischer Scientific Model FB505 sonicator, produced 300-600 bp fragments. Chromatin was clarified by centrifuging at 13,000 rpm for 15 min, diluted 1:1 with 2 X RIPA buffer (2 % Triton X-100, 0.2 % Na-deoxycholate, 0.2 % SDS, 280 mM NaCl, 20 mM Tris-HCl pH 8.0, 2 mM EDTA, 0.02 % Na-azide, 2 mM DMSF with complete protease inhibitor (Roche)). Chromatin solution (5.5 ml) was preabsorbed by incubation at 4° for 30 min with 55 μl of blocked Pierce™ Protein A agarose beads (Catalog #20333) and aliquots stored at -80°.

For larval chromatin, a modified protocol from (KUZU *et al.* 2016) was used. 150 larvae were frozen in liquid N_2_ and ground in a chilled mortar. The powder was transferred to a cooled 15 ml Dounce and homogenized with a loose pestle (10 strokes) and a tight pestle (15 strokes) in 10 ml PBS with protease inhibitor. Homogenate was made to 40 ml with PBS, crosslinked with 1 % formaldehyde for 20 min and quenched with 125 mM glycine for 30 mins. Crosslinked material was pelleted, washed once with wash buffer A (10 mM Hepes pH 7.6, 10 mM EDTA, 0.5 mM EGTA, 0.25 % Triton X-100, protease inhibitor and 0.2 mM PMSF), once with wash buffer B (10 mM Hepes pH 7.6, 100 mM NaCl, 1 mM EDTA, 0.5 mM EGTA, 1 % Triton X-100, protease inhibitor and 0.2 mM PMSF), and 3 times with TE wash buffer (10 mM Tris pH 8.0, 1 mM EDTA, 0.01 % SDS, protease inhibitor and 0.2 mM PMSF). The pellet was resuspended in 2 ml pre-RIPA buffer (0.1 % SDS, 10 mM Tris-HCl, 1 mM EDTA, protease inhibitor and 0.2 mM PMSF). Sonication was performed at settings described above for 2 min. Sonicated samples were diluted with 1 % Triton X-100, 0.1 % Na-deoxycholate, and 140 mM NaCl, centrifuged at 1500 g to clarify, aliquoted and stored at -80°.

### Chromatin Immunoprecipitation

Seventy five micrograms of chromatin was incubated overnight at 4° with 4 μl anti-H3K9me2 (Abcam, ab1220) or 8 μl anti-H3K9me3 (Abcam, ab8898), clarified by centrifugation at 14,000 rpm for 5 min and supernatants transferred to tubes containing 40 μl blocked Pierce™ Protein A agarose beads (Catalog #20333) and incubated 4 h at 4°. Following washing, reverse crosslinking, organic extraction and precipitation, DNA was suspended in 50 μl distilled water.

### ChIP-qPCR

Duplicate 20 μl reactions consisting of 2 μl DNA, 10 μl BioRad iTaq (#172-5101), and primers were amplified using an Mx3000P Real-Time PCR system (Stratagene).Standard error was derived from the mean Ct values of biological replicates. qPCR analysis was previously described (KOYA AND MELLER 2015). Each ChIP pull down was validated. For H3K9me2, primers in an H3K9me2-enriched region of the 3^rd^ chromosome and *dmn* served as positive and negative controls. ChIP primers are presented in Table S1. Primer specificity for 1.688^X^ repeats was ensured by anchoring one primer in flanking unique sequence (1.688^1A^, 1.688^3C^, 1.688^4A^ and 1.688^7E^) or by designing primers to unique sequences within repeats and testing with genomic DNA from a strain deleted for the repeat cluster (1.688^3F^ and 1.688^7F^; see Menon et al., 2014). Primer efficiencies were determined using MxPro qPCR software. Repeat copy number is normalized by expressing enrichment as percent input.

### Protein Isolation from embryos

Fifty mg of 0 - 12 hr embryos were homogenized in 250 μl RIPA buffer on ice. Homogenate was passed through a 26 gauge needle 10 - 12 times to shear DNA. Particulate matter was removed by centrifugation, and supernatant was mixed with an equal volume of 2 X SDS Sample buffer and boiled for 5 min before separation on a 15 % SDS polyacrylamide gel.

### Protein blotting

Polyacrylamide gels were equilibrated in transfer buffer (48 mM Tris, 39 mM glycine, 1.3 mM SDS, 20 % methanol) for 20 min. A PVDF membrane was activated in 100 % methanol for 1 min. Filter paper and activated PVDF membranes were saturated in transfer buffer and proteins transferred using a Trans-Blot SD Semi-Dry Transfer Cell (BIO-RAD). The membrane was washed in TBST (10 mM Tris-Cl, 200 mM NaCl, 0.1 % Tween 20, pH 7.5), blocked in 5 % BSA and probed overnight at 4° using 1:2000 mouse anti-H3K9me2 diluted in blocking solution (Abcam, ab1220) or 1:4000 goat anti-tubulin (Developmental Studies Hydrinoma Bank, E7). After washing with TBST, the membrane was incubated with alkaline phosphatase conjugated secondary antibodies (goat anti-mouse, Sigma, A3562 or rabbit anti-goat, Sigma, A4062), washed and developed in 100 mM diethanolamine, 100 mM NaCl, 5 mM MgCl_2_, pH 9.5 containing 33 μg/ml Nitroblue Tetrazolium (NBT) and 165 μg/ml 5-Bromo-4-chloro-3-indolyl phosphate (BCIP). Signals were quantified by ImageJ.

### Quantitative RT-PCR

Total RNA was isolated from 50 third instar male larvae or 100 mg dechorionated embryos using Trizol reagent (Invitrogen) as previously described (KOYA AND MELLER 2015). One microgram of RNA was reverse-transcribed using random hexamers and ImProm-II reverse transcriptase (Promega). Duplicate reactions were amplified using iTaq Universal SYBR Green Supermix (Bio-Rad) with an Mx3000P Real-Time PCR system (Stratagene). Primers are in Table S1. For determining relative transcript abundance, values were normalized to *dmn* (DCTN2-p50). To calculate fold change values were normalized to *dmn* and to a reference strain. Expression was calculated using the efficiency corrected comparative quantification method (PFAFFL 2001).

Strains and materials used in this study are available upon request. The authors affirm that all data necessary to confirm the conclusions of this study are presented within the article, figures and tables.

## Results

### Ago2 localizes at 1.688^X^ repeats

We took advantage of the resolution of ChIP and a FLAG-tagged *Ago2* transgene to determine if Ago2 localizes to 1.688^X^ repeats. FLAG-Ago2 was first tested for rescue of the dosage compensation function of Ago2. Males with the partial loss of function *roX1*^*ex40*^*roX2Δ* chromosome have high survival, as do *Ago2*^*-/-*^ flies, but synthetic lethality is observed in *roX1*^*ex40*^*roX2Δ*/Y; *Ago2*^*-/-*^ males (MENON AND MELLER 2012). One copy of a FLAG-*Ago2* transgene rescues these males, demonstrating that the FLAG tag does not disrupt the dosage compensation function of Ago2 (Figure 1A). Chromatin from FLAG-*Ago2*; *Ago*^*-/-*^ embryos, and from a reference strain lacking the FLAG-*Ago2* transgene, was immunoprecipitated with anti-FLAG antibodies and enrichment determined by quantitative PCR (qPCR). FLAG-Ago2 was enriched at the *Hsp70* promoter, a site known to bind Ago2 (CERNILOGAR *et al.* 2011) (Figure 1B). In contrast, a control region in the *dmn* gene displayed no enrichment. We then examined FLAG-Ago2 enrichment at a panel of six representative 1.688^X^ repeat clusters that differ in location and environment (within, near or far from protein coding genes), transcription level and sequence (Table 1). Interestingly, five of these show enrichment of FLAG-Ago2 over the repeats, but little or no enrichment in flanking regions (Figure 1C). We conclude that Ago2 localizes at many 1.688^X^ repeats, a finding that is consistent with involvement of Ago2 in siRNA-directed recruitment of chromatin modification at or around these regions.

**Figure 1.**
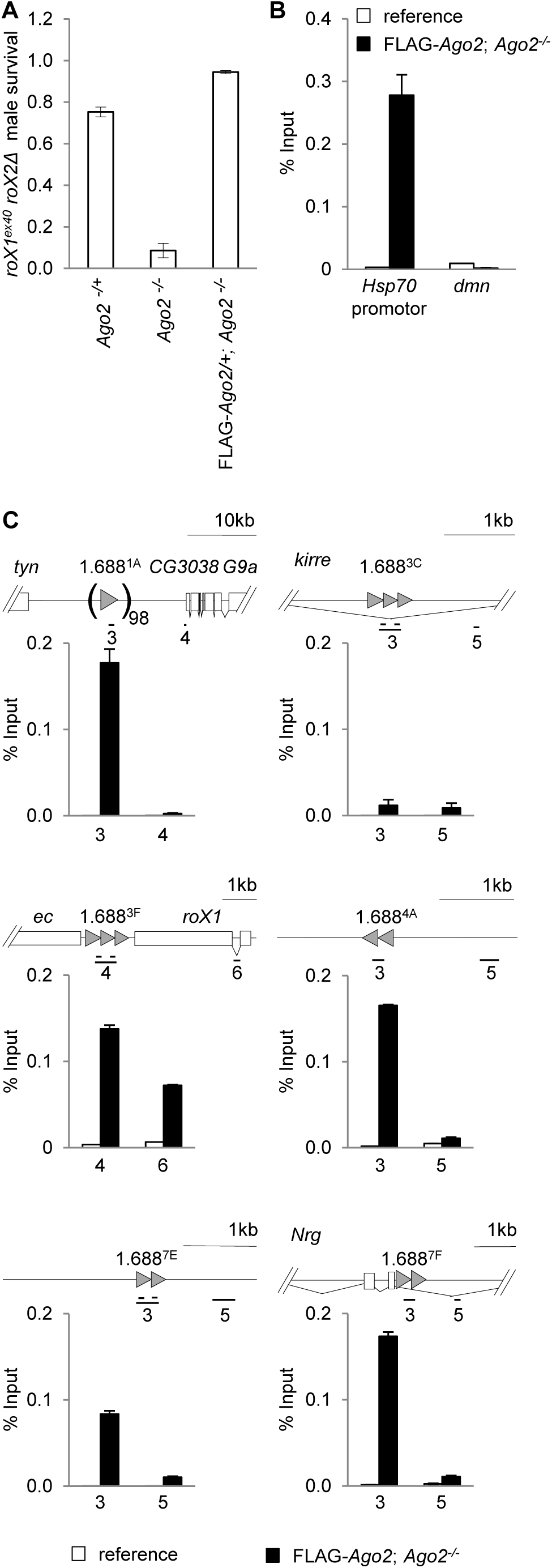
FLAG-Ago2 rescues the Ago2 dosage compensation function and localizes at 1.688^X^ repeats. (A) A FLAG-*Ago2* transgene (right) rescues the synthetic lethality of *roX1*^*ex40*^ *roX2Δ*/Y; *Ago2*^*414/414*^ males (center). (B) Chromatin from the laboratory reference strain (white) and *Ago2*^*414/414*^; FLAG*-Ago2* (black) embryos was precipitated with anti-FLAG antibody. Enrichment normalized to input is shown. The *Hsp70* promoter displays enrichment, but a control region in *dmn* does not. (C) FLAG-Ago2 enrichment is detected at several 1.688^X^ repeats (gray arrowheads).Approximately 100 copies of the 1.688^1A^ repeats are situated between *tyn* and CG3038. The 1.688^3C^ repeats are within a large *kirre* intron (splicing indicated by diagonal lines). Primers, indexed by gene and amplicon number, are presented in Table S1. Amplicon numbers, constant throughout this study, denote regions in selected repeats and flanking regions as indicated on gene models. In panel C only two amplicons per repeat, one including the repeats and in an adjacent region, were analyzed.

**Table I.** Panel of 1.688^X^ repeats used in this study. Cytological positions and scaffold coordinates were determined from Flybase (Release 6). The copy number at some positions differed between the laboratory reference strain and the genomic scaffold (see File S1). Similarity to 1.688^3F^ was determined by BLAST alignment of a 359 bp monomer. EST abundance is estimated from Flybase assignments.RNA poIII enrichment is derived from ChIP-seq of 6-8 h mesoderm (MONFORT *et al.* 2017). ^a^Quantitative RT-PCR (qRT PCR) is normalized to repeat copy number (see Figure S4).

| Repeat |  | 1.688<sup>1A</sup> | 1.688<sup>3C</sup> | 1.688<sup>3F</sup> | 1.688<sup>4A</sup> | 1.688<sup>7E</sup> | 1.688<sup>7F</sup> |
| --- | --- | --- | --- | --- | --- | --- | --- |
| Cytological position |  | 1A1 | 3B5 | 3F3 | 4A4 | 7E5 | 7F3 |
|  | Scaffold coordinates on X | 204,046..<br>241,257 | 2,768,669..<br>2,770,136 | 3,857,647..<br>3,858,186 | 4,070,631..<br>4,071,316 | 8,369,270..<br>8,369,783 | 8,530,749..<br>8,531,260 |
| Copy number | Genomic Scaffold | 98 | 3 | 2.5 | 2 | 1.5 | 1.5 |
|  | Lab reference | - | 3.5 | 3.5 | 2.5 | 2.5 | 1.5 |
| Repeat properties | Percent match to 1.688<sup>3F</sup> | 89 | 69 | 100 | 95 | 71 | 69 |
|  | Genomic environment | 37kb of repeats between convergent genes | Intronic | Between convergent genes | Isolated | Isolated | Intronic |
| Transcription | EST abundance | 0 | 2 | 9 | 2 | 0 | 26 |
|  | qRT PCR<sup>a</sup> | 0.00026 | 0.01 | 0.3 | 0.1 | 0.04 | 1.5 |
|  | RNA polII enrichment | - | - | + | - | - | - |

### Genetic interactions identify an Ago2-interaction network that participates in dosage compensation

Argonaute proteins in the RNA Induced Transcriptional Silencing (RITS) complexes of *S. pombe* and plants recruit chromatin modifiers to nascent transcripts (reviewed in (MELLER *et al.* 2015)). To explore the possibility of Ago2-interacting proteins participating in X chromosome recognition, we screened genes in an Ago2-interaction network for genetic interaction with *roX1 roX2*. A map of high probability Ago2-interactors was created using BioGRID (STARK *et al.* 2006), and esyN (BEAN *et al.* 2014) (Figure 2A; see File S2 for inclusion criteria). Members of this network were examined for genetic interactions with the partial loss of function *roX1*^*ex33*^*roX2Δ* X chromosome. *roX1*^*ex33*^*roX2Δ* males display partial mislocalization of MSL proteins and eclose at 20 % of normal levels (DENG *et al.* 2005b). Reduction of proteins that participate in X recognition further disrupts X localization and enhances *roX1*^*ex33*^*roX2Δ* male lethality (MENON AND MELLER 2012). Females are fully viable and fertile when the *roX* genes are mutated. *roX1*^*ex33*^*roX2Δ* females were mated to males that were heterozygous for a mutation in the gene being tested (Figure S1A). All sons are *roX1*^*ex33*^*roX2Δ*/Y, and heterozygous (experimental) or wild type (control) for the gene of interest. A reduction in normalized survival (experimental /control) reveals enhancement of *roX1 roX2* male lethality (Figure 2B, C). Daughters, which do not dosage compensate and are heterozygous for *roX1*^*ex33*^*roX2Δ,* do not display altered survival upon mutation of Ago2-interacting genes. As *G9a* is located on the X chromosome, a modified strategy to test this gene is presented in Figure S1B.

**Figure 2.**
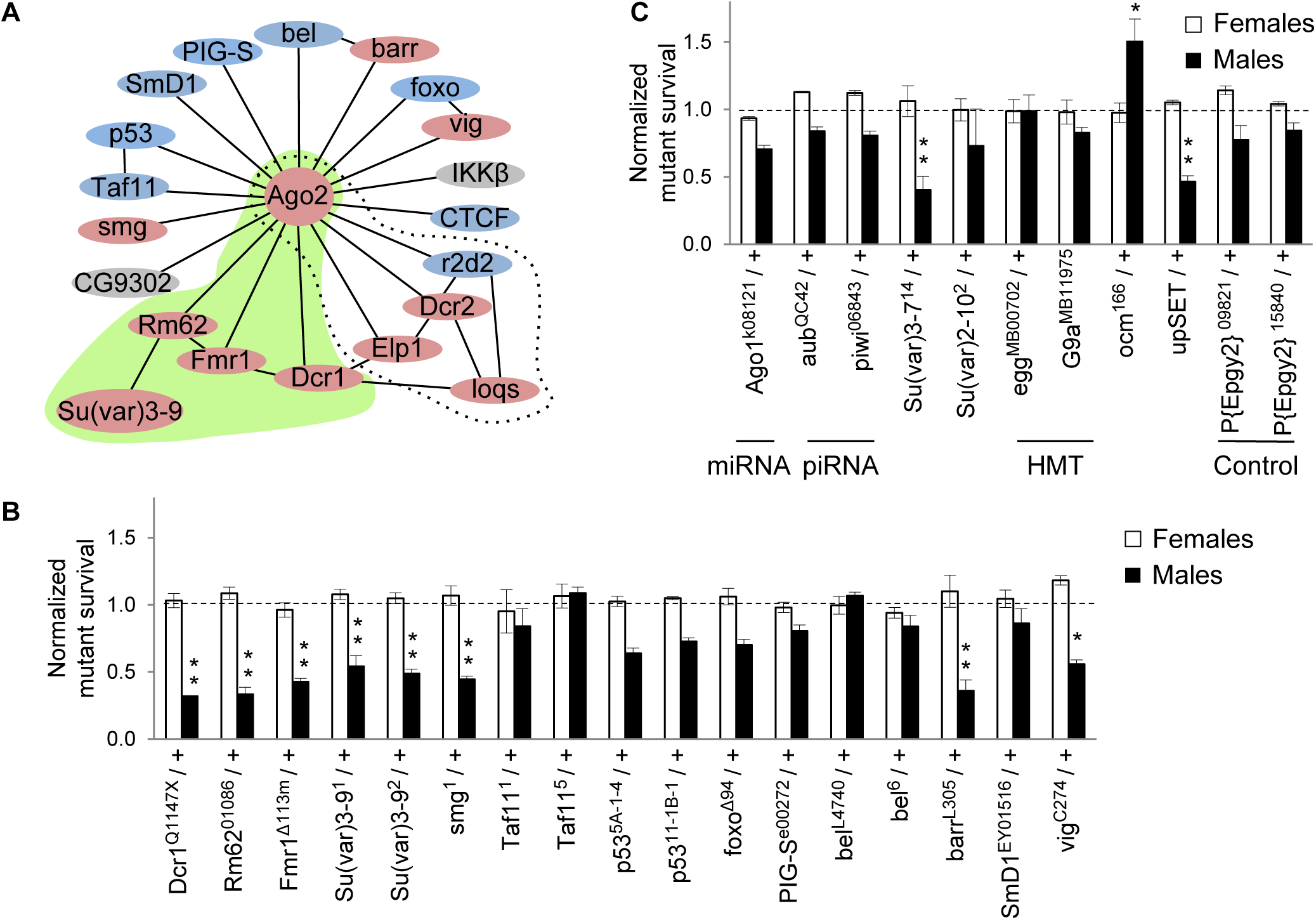
Ago2-interactors participate in dosage compensation. (A) Map of Ago2-interacting proteins. Genes displaying a genetic interaction with *roX1*^*ex33*^*roX2Δ* are pink, and those for which a significant interaction has not been detected are blue.Genes in gray are untested. A previously reported siRNA-producing sub-network is denoted by the dotted line. A putative chromatin-modifying sub-network identified in the present study is highlighted in green. Well-curated, high probability interactions from BioGRID and esyN are depicted by solid lines. See File S2 for inclusion criteria.(B) Mutations in many Ago2-interacting proteins reduce the recovery of *roX1*^*ex33*^*roX2Δ* males (black; *roX1*^*ex33*^*roX2Δ/Y; mut/+* normalized to *roX1*^*ex33*^*roX2Δ/Y; +/+*). Females are unaffected (white; *roX1*^*ex33*^*roX2Δ/++; mut/+* normalized to *roX1*^*ex33*^*roX2Δ/++; +/+*).(C) Additional controls and genes of interest. The mating strategy to test X-linked G9a is presented in Figure S1B. See Materials and Methods for *upSET* description. SEM is represented by error bars. Significance of ≤0.05 (*) and ≤0.001 (**) was determined using the Student’s two sample *t*-test.

Normalized survival of *roX1*^*ex33*^*roX2Δ* males with mutations in the Ago2-interaction network is presented in Figure 2B. Genes displaying significant interactions are noted by pink symbols, and those showing no interaction are blue in Figure 2A. We confirmed a previously identified siRNA-processing sub-network containing *Dcr2, Elp1*, and *loqs* (Figure 2A, dotted line; (MENON AND MELLER2012)). The present study identified several additional Ago2-interactors, including a potential chromatin-modifying sub-network containing *Dcr1, Fmr1, Rm62,* and the histone methyltransferase *Su(var)3-9* (green, Figure 2A). Su(var)3-9 deposits H3K9me2 and acts with Rm62 to re-silence active chromatin (BOEKE *et al.* 2011).

Additional chromatin modifiers and genes in other small RNA pathways were also tested (Figure 2C). A previous study found no interaction between *roX1*^*ex33*^*roX2Δ* and the piRNA pathway genes *aub* and *piwi,* or the miRNA pathway gene *Ago1*, a finding replicated here (MENON AND MELLER 2012). Since our findings point towards involvement of chromatin modifiers, we tested the chromatin regulatory factor *Su(var)2-10* and two additional H3K9 methyltransferases, *eggless* (*egg*) and *G9a* (Figure 2C).None of these modified *roX1*^*ex33*^*roX2Δ* survival Mutations in *Su(var)3-7*, important for heterochromatin formation, and *upSET*, which maintains heterochromatin and H3K9me2 levels, enhance *roX1*^*ex33*^*roX2Δ* male lethality (SPIERER *et al.* 2008; MCELROY *et al.* 2017). *Over compensating males* (*ocm)* has an unusual dosage compensation phenotype as mutations in *ocm* rescue males with insufficient MSL activity, suggesting that it might act as a governor of activation (LIM AND KELLEY 2013). Interestingly, mutation of *ocm* significantly increased the survival of *roX1*^*ex33*^*roX2Δ* males, supporting the idea that *ocm* normally restrains activation. The P{EPgy2}^*09821*^ and P{EPgy2}^15840^ strains, used to outcross *Su(var)3-9* and *barr* mutants, display no interaction and serve as controls for genetic background. Taken together, these findings suggest that several genes that deposit H3K9me2, maintain this mark or participate in heterochromatin formation also contribute to X chromosome dosage compensation. At first glance these observations appear to be at odds with X chromosome hypertranscription, the ultimate consequence of X chromosome recognition.

### Ectopically expressed 1.688^3F^ siRNA disrupts H3K9me2 patterns

Previous studies found that ectopically produced 1.688^3F^ siRNA partially rescues *roX1 roX2* males and increases X localization of the MSL complex (MENON *et al.* 2014). The mechanism by which siRNA promotes X recognition is unknown. The discovery that insertion of 1.688^X^ DNA on an autosome enables functional compensation of nearby genes, and the enhancement of this effect by ectopic 1.688^3F^ siRNA, suggests siRNA action through cognate genomic regions (JOSHI AND MELLER 2017). In accord with this idea, an autosomal *roX1* transgene also enables compensation of nearby genes, but is unaffected by 1.688^3F^ siRNA. To test the idea that 1.688^3F^ siRNA directs epigenetic modification of 1.688^X^ chromatin, we used ChIP to analyze chromatin around 1.688^X^ repeats on the X chromosome. ChIP-qPCR detected H3K9me2 enrichment in 4 out of 6 repeats (white bars, Figure 3). As H3K9me2 enrichment was not uniform, we considered additional factors that might determine this mark, and noted that only repeats showing evidence of transcription were enriched for H3K9me2, consistent with the idea of Ago2-dependent recruitment to nascent transcripts (Figure S4) (VERDEL *et al.* 2004). Upon ectopic expression of 1.688^3F^ siRNA a dramatic disruption of H3K9me2 was observed in and around 1.688^X^ repeats (black bars, Figure 3). For example, 1.688^3F^ and 1.688^4A^ display peaks of H3K9me2 in wild type flies, but this mark was reduced over the repeats and increased in surrounding regions by elevated 1.688^3F^ siRNA. The reduction in H3K9me2 over repeats themselves was unexpected and could represent repositioning of nucleosomes or blocking of the methylated site by another protein. Untranscribed repeat clusters at 1.688^1A^ and 1.688^7E^ show no H3K9me2 enrichment in wild type flies, but gained H3K9me2 upon expression of 1.688^3F^ siRNA. In contrast, no enrichment of H3K9me3 in or near 1.688^X^ repeats was detected in wild type or 1.688^3F^ siRNA-expressing embryos (Figure S2). We conclude that some 1.688^X^ repeats are enriched for H3K9me2, and that cognate siRNA broadly disrupts this mark within and several kb adjacent to 1.688^X^ DNA.

**Figure 3.**
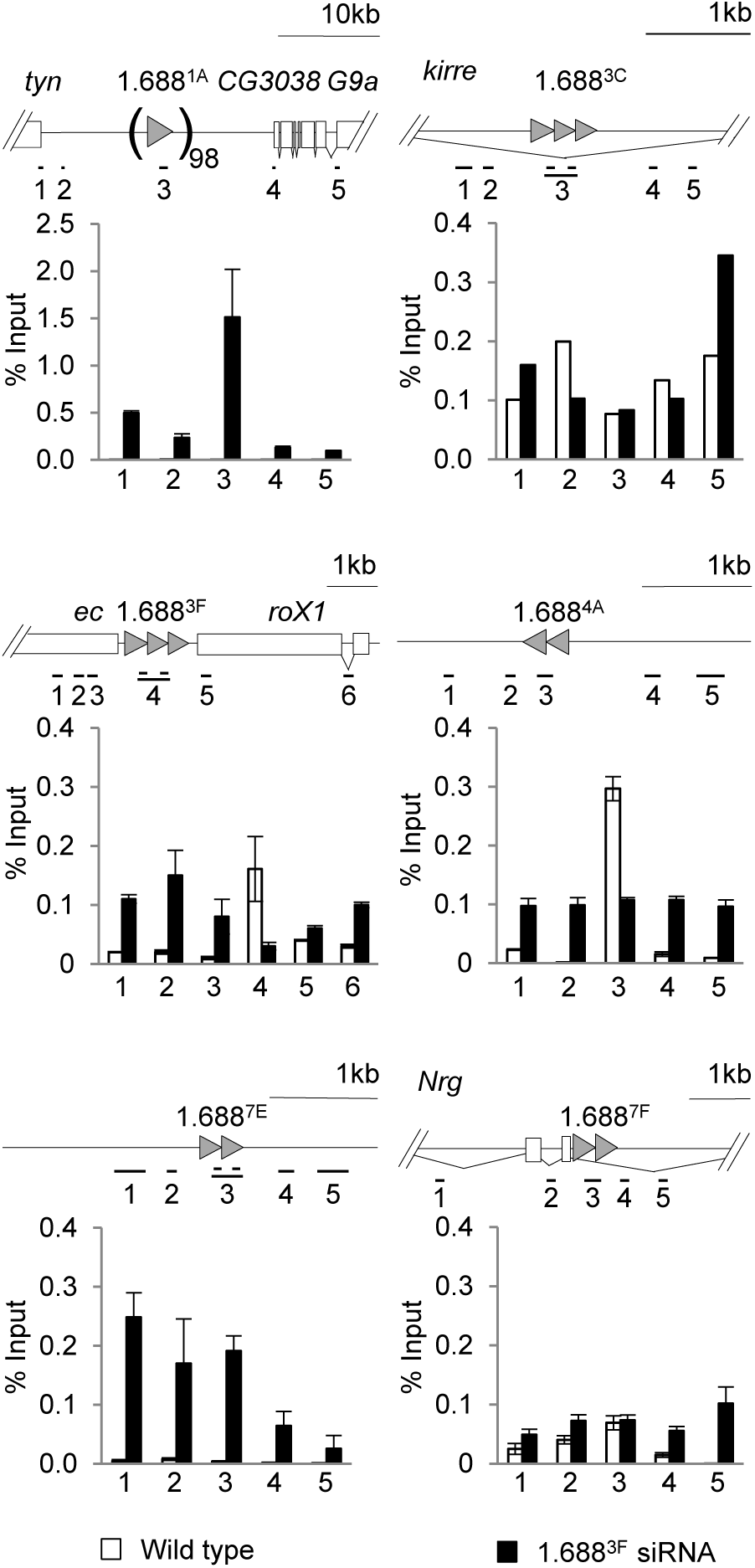
Elevated 1.688^3F^ siRNA disrupts H3K9me2 enrichment around 1.688^X^ repeats. Chromatin from wild type embryos (white) and embryos ectopically producing 1.688^3F^ siRNA (black) was immunoprecipitated with antibody to H3K9me2. Enrichment over input was determined by quantitative PCR (qPCR). The standard error of two biological replicates is shown. Amplicons correspond to numbered positions on the gene models above. Primers are presented in Table S1.

### Su(var)3-9 deposits H3K9me2 at 1.688^X^ repeats

The identification of Su(var)3-9 as an indirect binding partner of Ago2, observation of a genetic interaction between *roX1 roX2* and *Su(var)3-9* and enrichment of H3K9me2 on some 1.688^X^ repeats suggested that Su(var)3-9 could be modifying 1.688^X^ repeats. *D. melanogaster* has 3 histone H3K9 methyltransferases, *Su(var)3-9, eggless*, and *G9a*, but only *Su(var)3-9* mutations enhance the male lethality of *roX1 roX2* (Figure 2; (SWAMINATHAN *et al.* 2012)). To determine if Su(var)3-9 is responsible for H3K9me2 at 1.688^X^ chromatin, we generated strains carrying *Su(var)3-9* over a marked balancer, enabling selection of homozygous *Su(var)3-9* mutant larvae. H3K9me2 enrichment is virtually eliminated over 1.688^X^ repeats in *Su(var)3-9*^-/-^ mutants (Figure 4, gray) and remains low in *Su(var)3-9*^*-/-*^ larvae that express 1.688^3F^ siRNA (Figure 4, black). This reveals that Su(var)3-9 deposits H3K9me2 at 1.688^X^ chromatin in wild type flies, and eliminates the possibility that a different methyltransferase is recruited to these regions following ectopic expression of 1.688^3F^ siRNA. Disruption of H3K9me2 upon expression of 1.688^3F^ siRNA thus reflects changes in the localization or activity of Su(var)3-9.

**Figure 4.**
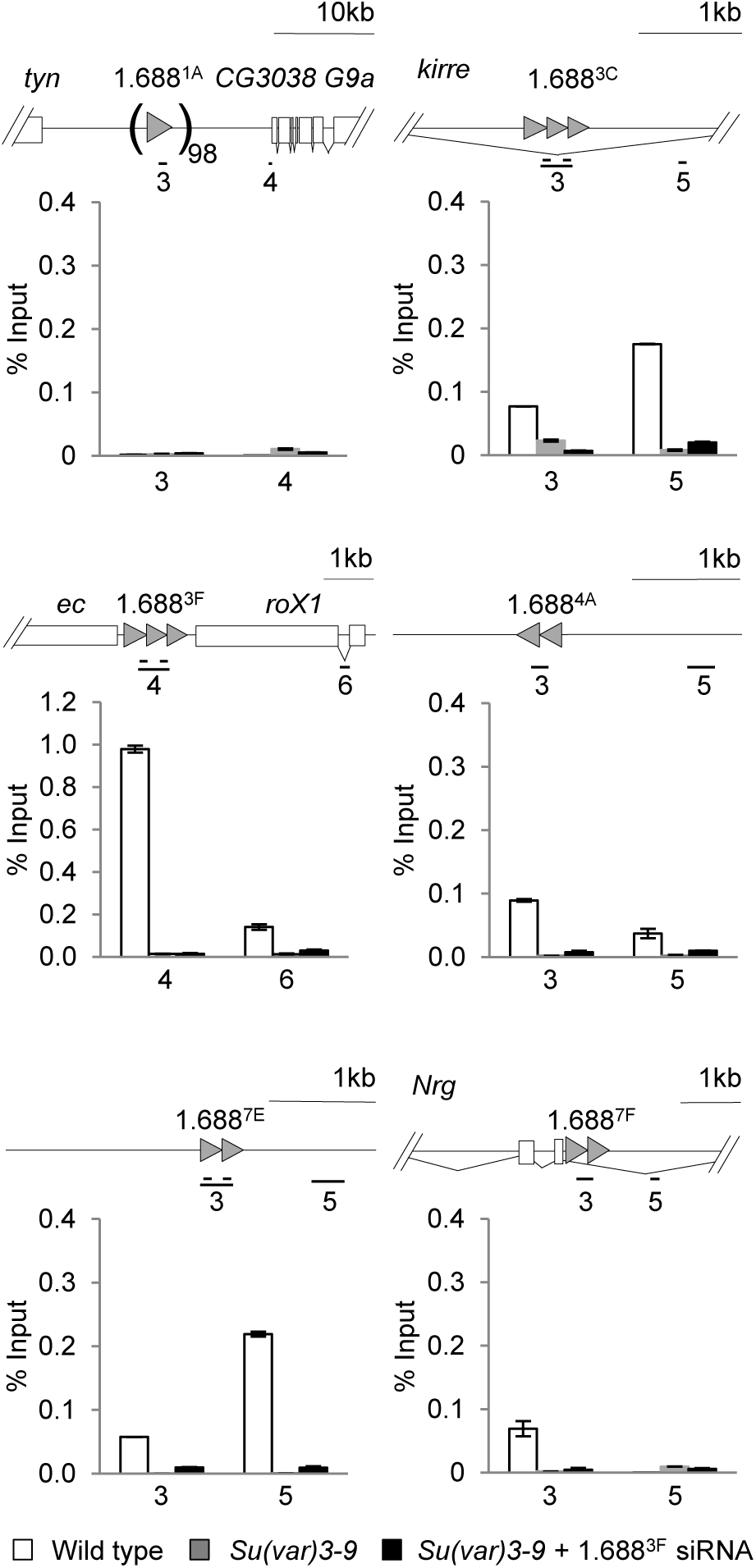
Su(var)3-9 deposits H3K9me2 at 1.688^X^ repeats. Chromatin from wild type male larvae (white), *Su(var)3-9*^*1*^/*Su(var)3-9*^*1*^ male larvae (gray), and *Su(var)3-9*^*1*^*/Su(var)3-9*^*1*^ males ectopically expressing 1.688^3F^ siRNA (black) was immunoprecipitated with antibody to H3K9me2. Enrichment normalized to input is shown. Standard error is derived from two biological replicates. See materials and methods for full genotypes and larval selection strategy.

To determine how far from 1.688^X^ repeats the H3K9me2 disruption extends, regions 10 to 26 kb from repeats were examined. In each case, increased H3K9me2 was observed in embryos with ectopic 1.688^3F^ siRNA expression (Figure S3A). This suggested the possibility of a global change in H3K9me2 levels. To address this we probed protein blots from wild type and 1.688^3F^ siRNA-expressing embryos to determine global levels of this modification, but found no evidence of a change in H3K9me2 (Figure S3B). As most H3K9me2 is found in heterochromatic regions that comprise > 30 % of the fly genome, changes in euchromatin may represent a negligible portion of the nuclear pool.

H3K9me2 is generally thought to be repressive, but compensation in flies occurs by increased expression of X-linked genes. To determine if changes in H3K9me2 enrichment correlate with changes in transcription, regions flanking 1.688^X^ repeats were examined in wild type and 1.688^3F^ siRNA-expressing embryos. Consistent with H3K9me2 having a repressive effect, 1.688^3F^ siRNA decreases accumulation of mRNA, as well as non-coding intragenic and intronic regions, with elevated H3K9me2 (Figure S4). The apparent increase in 1.688^3F^ expression presumably originates from the transgene used to produce ectopic 1.688^3F^ siRNA. We detected dramatic reductions in messages immediately adjacent to 1.688^1A^ (*tyn, G9a*) and 1.688^3F^ (*ec, roX1*). In spite of a 90 % reduction in *ec* transcript in embryos expressing 1.688^3F^ siRNA, adults do not display the rough eye *ec* phenotype. It is possible that ectopic 1.688^3F^ siRNA has a more pronounced effect in embryos, whose lack of differentiation may make them particularly susceptible to epigenetic disruption. Mature patterns of chromatin organization are established later in development, and these may be more resilient. To test this, we examined expression in 1.688^3F^ siRNA-expressing third instar male larvae, and found that *tyn, G9a* and *ec* regained wild type transcript levels, and *roX1* was largely restored (Figure S4). The precise reason for the differences between embryos and larvae are uncertain, but restoration of normal gene expression by the third larval instar is consistent with the lack of phenotype in otherwise wild type flies that ectopically express 1.688^3F^ siRNA (MENON *et al.* 2014).

The finding that animal age influenced response to ectopic siRNA prompted us to determine the time point at which H3K9me2 is established at 1.688^X^ repeats. A possible scenario is that this mark is placed before MSL localization, and acts in some way to guide X recognition. X-localization of the MSL complex occurs 3 h after egg laying (AEL) (RASTELLI *et al.* 1995; MELLER 2003). We measured H3K9me2 enrichment at 1.688^3F^ in embryos before the MSL complex binds to the X (1.5 - 3 h), during initial MSL recruitment (3 - 4 h), and at 4 - 6 h and 6 -12 h. In contrast to our prediction, H3K9me2 is first detected on 1.688^3F^ between 6 and 12 h AEL, after X localization of the MSL complex has occurred (Figure S5). We conclude that H3K9me2 at 1.688^X^ repeats is unlikely to guide initial X recognition, but may serve at a later time point to facilitate spreading of this mark or enforce the stability of X recognition. As lethality due to failure of dosage compensation occurs at the end of the 3^rd^ instar, rescue of male viability could presumably occur during a window of several days.

### H3K9me2 is enriched at regions flanking autosomal 1.688^3F^ transgenes

One challenge of studying recruiting elements on the X chromosome is that the redundancy and proximity of these elements complicates interpretation. To overcome this we tested autosomal integrations of 1.688^3F^ DNA or *roX1* (Figure 5A) (JOSHI AND MELLER2017). H3K9me2 ChIP was performed on chromatin from third instar male larvae with 1.688^3F^ (Figure 5B) or *roX1* (Figure 5C) on 2L (gray bars), and in the same genotypes with ectopic expression of 1.688^3F^ siRNA (black bars). H3K9me2 within 5 kb of the integration site is not strongly enriched in control males, or in males with a *roX1* transgene, but is striking elevated when 1.688^3F^ DNA is present. Consistent with our observations in embryos, ectopic 1.688^3F^ siRNA further elevated H3K9me2 near the 1.688^3F^ integration. This contrasts with negligible enrichment flanking the *roX1* transgene (Figure 5C). For unknown reasons, enrichment over the integrated 1.688^3F^ DNA was itself undetectable. We conclude that autosomal insertion of 1.688^3F^ DNA makes flanking chromatin subject to siRNA-induced H3K9me2 deposition. Taken together, these studies support the idea that the 1.688^X^ repeats influence patterns of H3K9me2 nearby, but CES-containing *roX1,* with a different class of recruiting element, has little effect.

**Figure 5.**
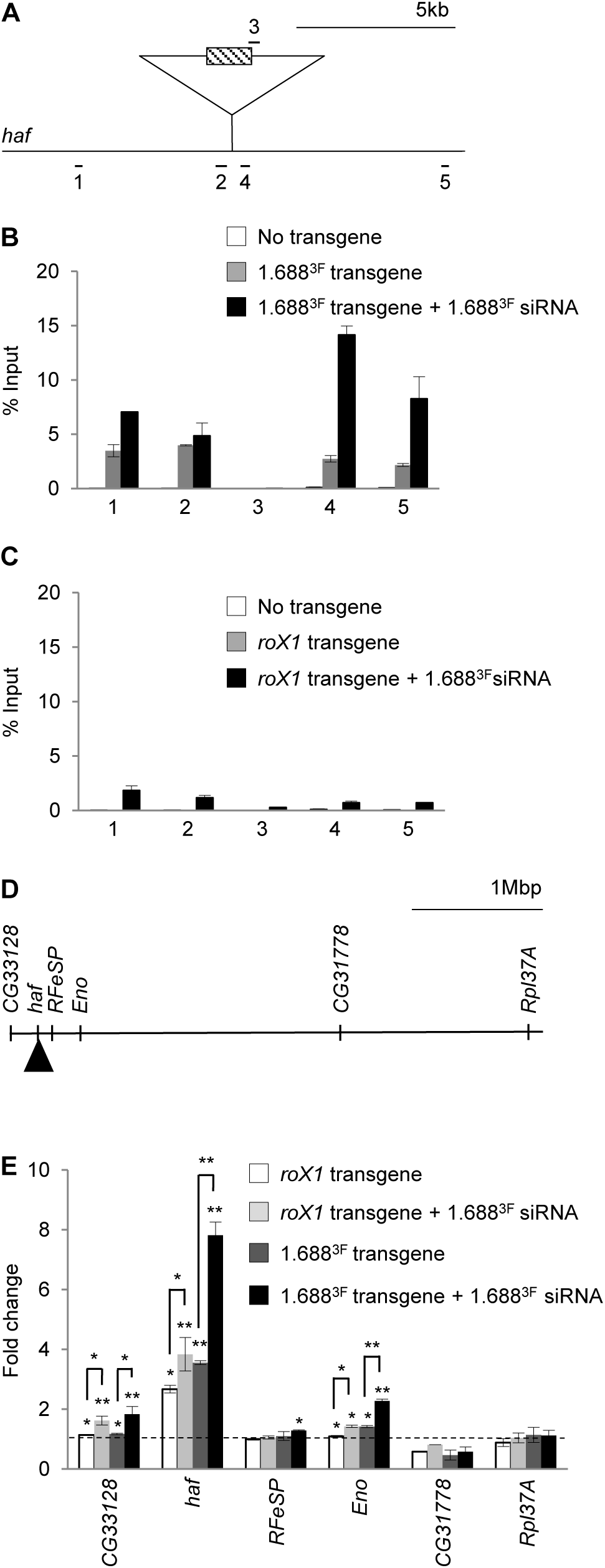
Ectopic 1.688^3F^ siRNA increases H3K9me2 flanking an autosomal 1.688^3F^ DNA insertion and elevates expression of nearby genes. (A) Amplicons flanking the landing site in a large *haf* intron at 22A3 (splicing not shown). (B) H3K9me2 enrichment surrounding the 1.688^3F^ transgene. Chromatin from wild type third instar male larvae (white), larvae with 1.688^3F^ DNA at the landing site (gray), and larvae with 1.688^3F^ DNA at the landing site and ectopic 1.688^3F^ siRNA (black) was immunoprecipitated with antibody to H3K9me2. (C) H3K9me2 enrichment surrounding a *roX1* insertion. Chromatin from wild type male third instar larvae (white), larvae with the *roX1* insertion (gray), and with the *roX1* insertion and ectopic 1.688^3F^ siRNA (black) was immunoprecipitated. Data is from two biological replicates and enrichment is normalized to input. (D) Portion of 2L showing relative location of *CG33128, haf, RFeSP, Eno, CG31778*, and *Rpl37A*. (E) Accumulation of transcripts in male larvae carrying *roX1* (white) or 1.688^3F^ insertions (dark gray), and in male larvae that express ectopic 1.688^3F^ siRNA and have *roX1* (light gray) or 1.688^3F^ integrations (black).Expression is normalized to *dmn* and wild type male larvae. SEM is derived from three biological replicates. Significance was determined using Student’s two sample *t*-test,≤0.05 (*), ≤0.001 (**) significance. Primers are presented in Table S1.

To determine the influence of 1.688^3F^ and *roX1* on transcription of autosomal genes on 2L, we performed quantitative RT-PCR (qRT PCR) on total RNA from third instar male larvae of the genotypes described above. The 1.688^3F^ and *roX1* integration site is in an intron of *haf,* over 17 kb from the closest exon. We also examined *RFeSP, CG33128, Eno* (30, 89 and 142 kb from the insertion, respectively), and *CG31778* and *Rpl37A*, 2.1 and 3.5 Mb distant (Figure 5D). The presence of 1.688^3F^ or *roX1* integrations alone had no effect on the most distant genes, *CG31778* and *Rpl37A*. A *roX1* integration increased expression of *haf* 2.5 fold, more than expected from full compensation. This may reflect the fact that MSL recruitment to a *roX1* transgene can overcome local, chromatin-based silencing (KELLEY AND KURODA 2003). Addition of 1.688^3F^ siRNA increased *haf* expression slightly, and similarly increased expression of *CG33128* and *Eno* (light gray bars, Figure 5E).

A 1.688^3F^ insertion produced a four-fold increase in *haf*, and a slight increase in *Eno*, 141 kb from the integration site. But upon expression of 1.688^3F^ siRNA, *haf* expression increased to 8 fold wild type levels, and *CG33128* and *Eno* both increased to 2-fold wild type levels, consistent with full compensation. We conclude that an autosomal insertion of 1.688^X^ DNA induces H3K9me2 deposition on flanking chromatin, but also increases expression of genes on 2L in a manner that is consistent with recruitment of the MSL complex. Both H3K9me2 enrichment and increased expression is enhanced by 1.688^3F^ siRNA, suggesting that X identification involves an siRNA-directed mechanism that operates through 1.688^X^ repeats.

## Discussion

Molecularly distinct dosage compensation strategies have arisen independently in different organisms, but a shared feature is the ability to selectively recognize and alter an entire chromosome. How a regulatory system is directed to a single chromosome is poorly understood. The discovery that 1.688^X^ satellite DNA promotes recruitment of dosage compensation to nearby genes supports the idea that these repeats are important for selective recognition of X chromatin (JOSHI AND MELLER 2017). How the 1.688^X^ repeats accomplish this is a question of great interest. Involvement of the siRNA pathway, and siRNA from a 1.688^X^ repeat, in X recognition points to the possibility that siRNA-directed modification of chromatin around 1.688^X^ repeats plays a role in dosage compensation in normal males. The findings of the current study support this idea.

Although numerous studies point to small RNA regulation of chromatin in flies, this process is better understood in other organisms (reviewed in (MELLER *et al.* 2015)). Small RNA directed heterochromatin formation was discovered in *S. pombe* (reviewed in (MOAZED 2009)). Heterochromatic regions are transcribed during S phase, and transcripts are processed into siRNAs that guide the Ago1-containing RITS complex to complementary, nascent transcripts (VERDEL *et al.* 2004). In addition to several other activities, RITS recruits the H3K9 methyltransferase Clr4 (ZHANG *et al.* 2008). We propose that a similar process is occurring at 1.688^X^ chromatin in flies. Most 1.688^X^ repeats bind Ago2, and many are transcribed. Several of the 1.688^X^ repeats that we examined are enriched for H3K9me2 deposited by Su(var)3-9, an ortholog of Clr4. Our screen identified genetic interactions between *roX1 roX2* and members of a possible RITS-like complex consisting of Ago2, Rm62 and Su(var)3-9. Finally, H3K9me2 enrichment in, and around, 1.688^X^ repeats is responsive to 1.688^X^ siRNA, and enrichment is blocked by loss of Su(var)3-9. Taken together, these findings suggest that a RITS-like complex normally modifies chromatin at 1.688^X^ repeats.

The idea that repressive H3K9me2 marks participate in a process culminating in a two-fold increase in expression is counterintuitive, but X recognition is complex. This process involves CES sites that directly recruit the MSL complex and the 1.688^X^ repeats, acting indirectly to enhance X recognition (Figure 6). It is possible that X recognition uses epigenetic marks, such as H3K9me2, that are distinct from the activating marks deposited within genes by the MSL complex. We propose that robust X recognition results from cooperation between two distinct pathways that guide this process. Interestingly, numerous studies have discovered links between the compensated X chromosome of male flies and repressive marks. For example, the male X is enriched in HP1, a major constituent of heterochromatin that binds H3K9me2 (DE WIT *et al.* 2005; LIU *et al.* 2005). The structure of the polytenized male X chromosome is extraordinarily sensitive to altered levels of genes that participate in heterochromatin formation or silencing, such as *HP1, Su(var)3-7* and *ISWI*. Mutations in these genes produce a general disruption of polytenization that is strikingly specific to the male X (DEURING *et al.* 2000; SPIERER *et al.* 2005; ZHANG *et al.* 2006). JIL-1, a kinase that enforces boundaries between heterochromatin and euchromatin, is enriched on the X chromosome and thought to participate in compensation (JIN *et al.* 2000; WANG *et al.* 2001; EBERT *et al.* 2004; DENG *et al.* 2005a). Upon loss of JIL-1, polytenized structure is disrupted and H3K9me2 invades euchromatic chromosome arms, but the X chromosome is most severely affected (ZHANG *et al.* 2006). Finally, the MSL proteins themselves have an affinity for heterochromatin. In *roX1 roX2* mutant males the MSL proteins become mislocalized to ectopic autosomal sites (MELLER AND RATTNER 2002).For reasons that are still unclear, the most prominent of these sites are the heterochromatic 4^th^ chromosome and chromocenter (DENG AND MELLER 2006; FIGUEIREDO *et al.* 2014). Taken together, these observations suggest that X recognition, or spreading of the MSL complex, could be facilitated by repressive marks. One intriguing possibility is that 1.688^X^ repeats guide deposition of H3K9me2 and this mark, directly or indirectly, assists localization of the MSL complex. Although MSL-mediated compensation initiates at 3 h AEL, before H3K9me2 enrichment over 1.688^X^ chromatin, male killing due to loss or mislocalization of the MSL complex occurs several days later at the larval/pupal transition. It is possible that ectopic production of 1.688^X^ siRNA drives enrichment of H3K9me2 across the X, supporting X recognition or MSL complex spreading at later developmental stages. This would explain why *roX1 roX2* mutant male larvae, defective for X recognition, display increased X-localization of the MSL proteins and elevated viability upon expression of 1.688^X^ siRNA (MENON *et al.* 2014).

**Figure 6.**
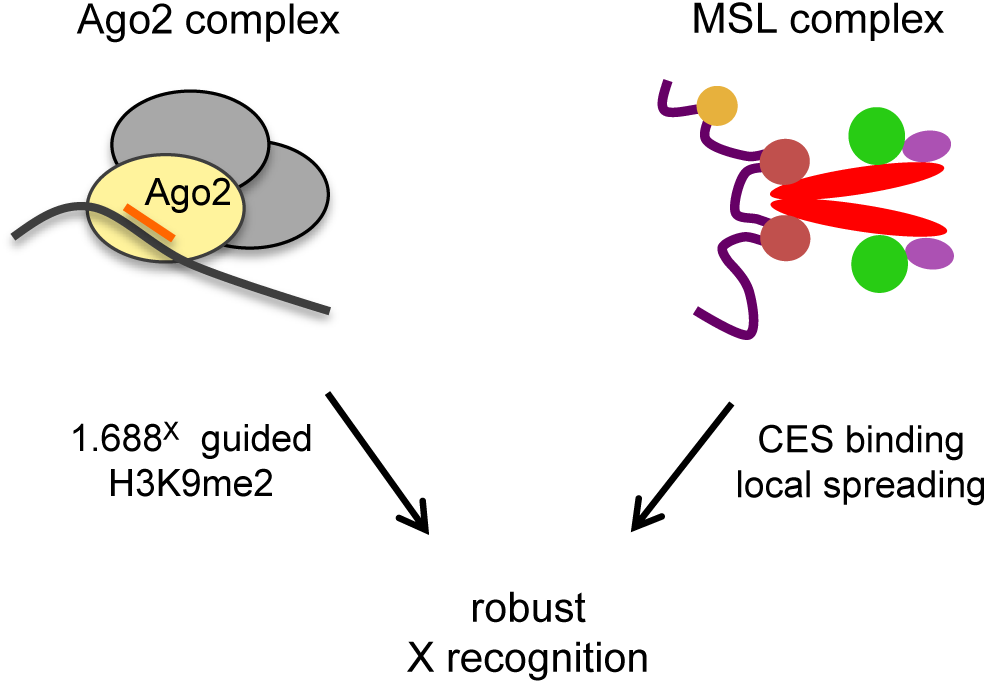
Proposed model of cooperative X recognition. The MSL complex is directly recruited to CES and then spreads into active genes nearby (right). 1.688^X^ siRNA guides an Ago2-containing complex that modifies chromatin around 1.688^X^ DNA, possibly in a transcription-dependent manner (left). We postulate that robust X recognition involves the cooperative action of both pathways.

An intriguing aspect of dosage compensation is the evolutionary convergence of mechanisms. For example, long non-coding RNA also plays a central role in X recognition in mammals, where expression of the *X inactive specific transcript* (*Xist*) RNA guides X inactivation (LEE 2009). Furthermore, repetitive LINE-1 elements that are enriched on the mammalian X chromosome are proposed to facilitate X inactivation (LYON 1998; BAILEY *et al.* 2000). Interestingly, some LINE-1 elements are transcribed during the onset of X inactivation, producing endo-siRNAs that may guide local spreading of heterochromatin into regions that are otherwise prone to escape (CHOW *et al.* 2010). These parallels are particularly striking as the outcomes, silencing of an X chromosome in mammalian females and activation of the single X in male flies, appear unrelated.We propose that cooperation between distinct chromatin-modifying systems that rely on long and short non-coding RNAs is one strategy to selectively modulate an entire chromosome.

## Acknowledgments

The authors wish to thank colleagues who donated fly strains, Drs. S. Todi, M.Kuroda,R. Kelley and P. Spierer. We thank Drs. S. K. Koya and E. Larschan for advice on ChIP. Stocks obtained from the Bloomington Drosophila Stock Center (NIH P40OD018537) and the Kyoto Stock Center were used in this study. N. D. was supported in part by a Summer Dissertation Fellowship and a Graduate Enhancement award. This work was supported by NIH grant GM 093110 to V.H.M.

---

**Figure S1.**
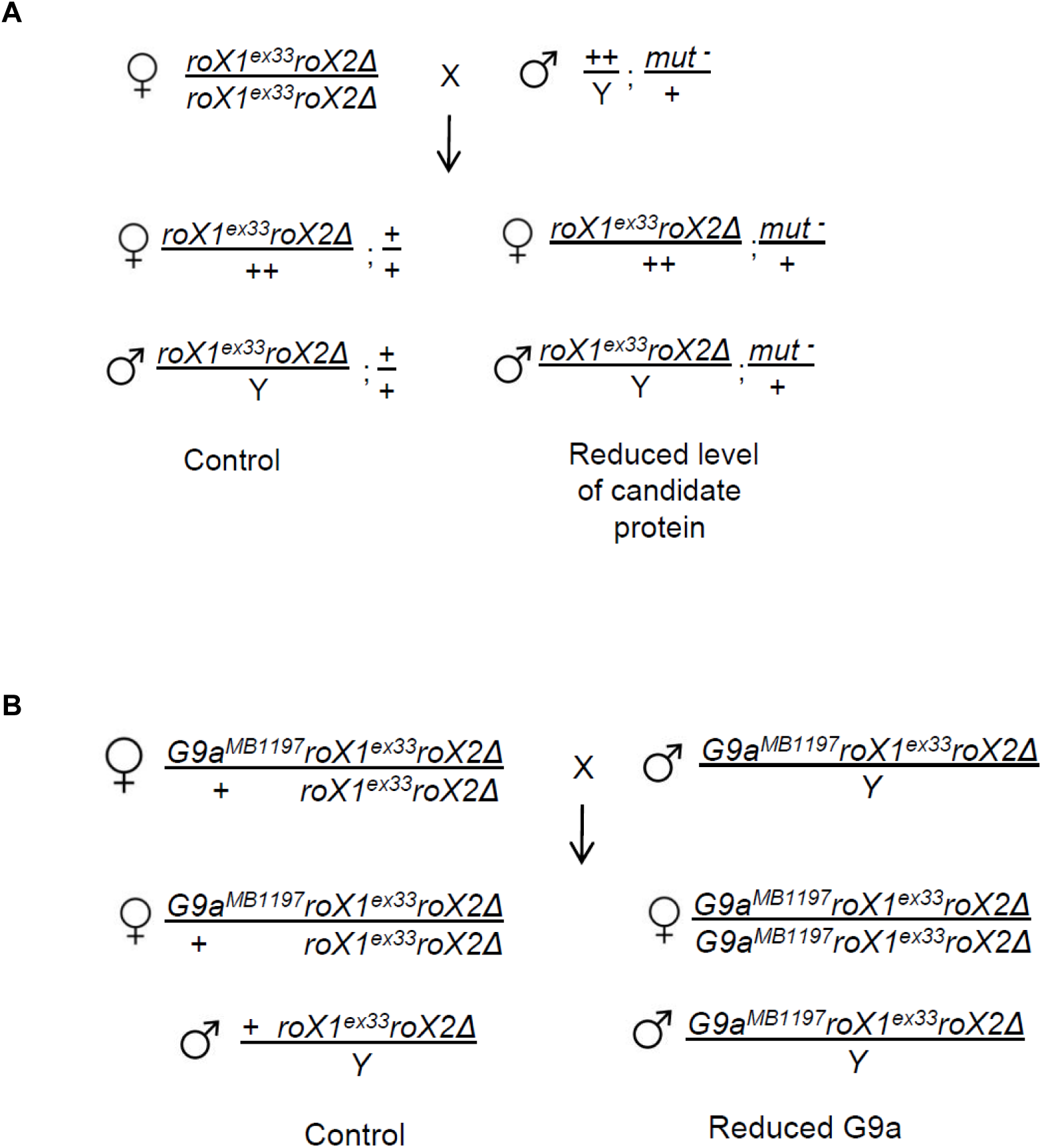
Matings to detect genetic interactions between *roX1 roX2* and candidate genes. (A) *roX1*^*ex33*^*roX2Δ* females were mated to males heterozygous for a mutation in the gene of interest. The survival of sons mutated for the gene of interest (bottom right) is divided by that of control brothers (bottom left) and presented in Figure 1B and C. In an otherwise wild type background, *roX1*^*ex33*^*roX2Δ* allows 20 % adult male escapers. Females do not dosage compensate and serve as an internal control.(B) Testing for genetic interaction between G9a and *roX1*^*ex33*^*roX2Δ*. Heterozygous *G9a*^*MB1197*^ *roX1*^*ex33*^*roX2Δ*/+ *roX1*^*ex33*^*roX2Δ* females were mated to *G9a* ^*MB1197*^ *roX1 roX2* males. *G9a*^*MB1197*^ is marked with EGFP. The survival of *G9a* ^*MB1197*^ *roX1*^*ex33*^*roX2Δ* sons (EGFP-positive, right) was divided by that of EGFP-negative + *roX1*^*ex33*^*roX2Δ* sons (left). EGFP intensity differentiates daughters that are homozygous or heterozygous for *G9a* ^*MB1197*^

**Figure S2.**
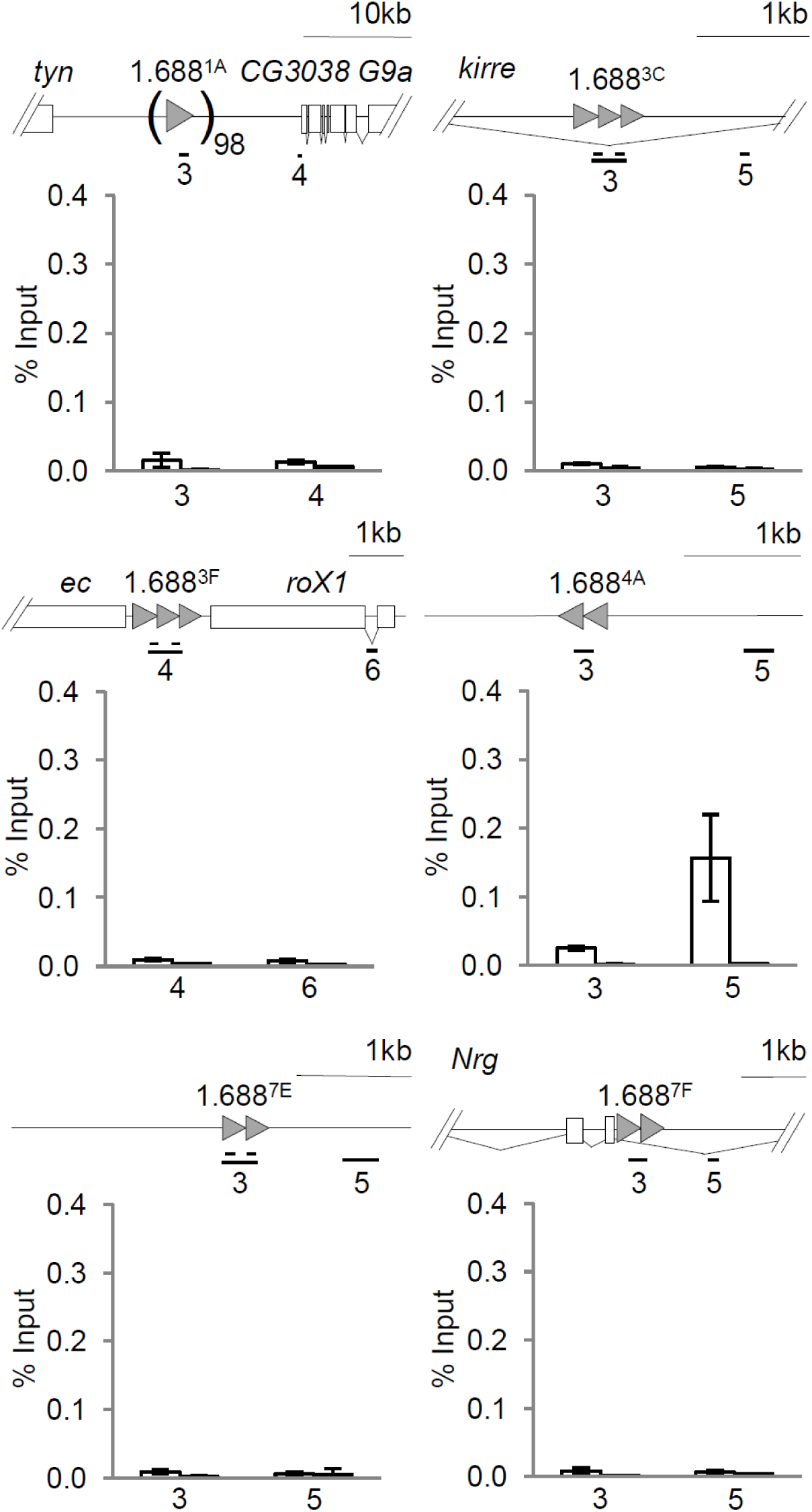
H3K9me3 is not enriched over 1.688^X^ repeats or altered by ectopic expression of 1.688^3F^ siRNA. Chromatin from 6-12 hr embryos was immunoprecipitated with antibody to H3K9me3. DNA was analyzed by qPCR using primers within 1.688^X^ repeats (gray triangles) or in flanking regions. Approximately 100 copies of 1.688^1A^ are present between *tyn* and *CG3038*. Primers indexed by gene and amplicon number are presented in Table S1. No significant enrichment within repeats, or change in H3K9me3 within repeats, is observed following siRNA expression. Standard error is derived from two biological replicates.

**Figure S3.**
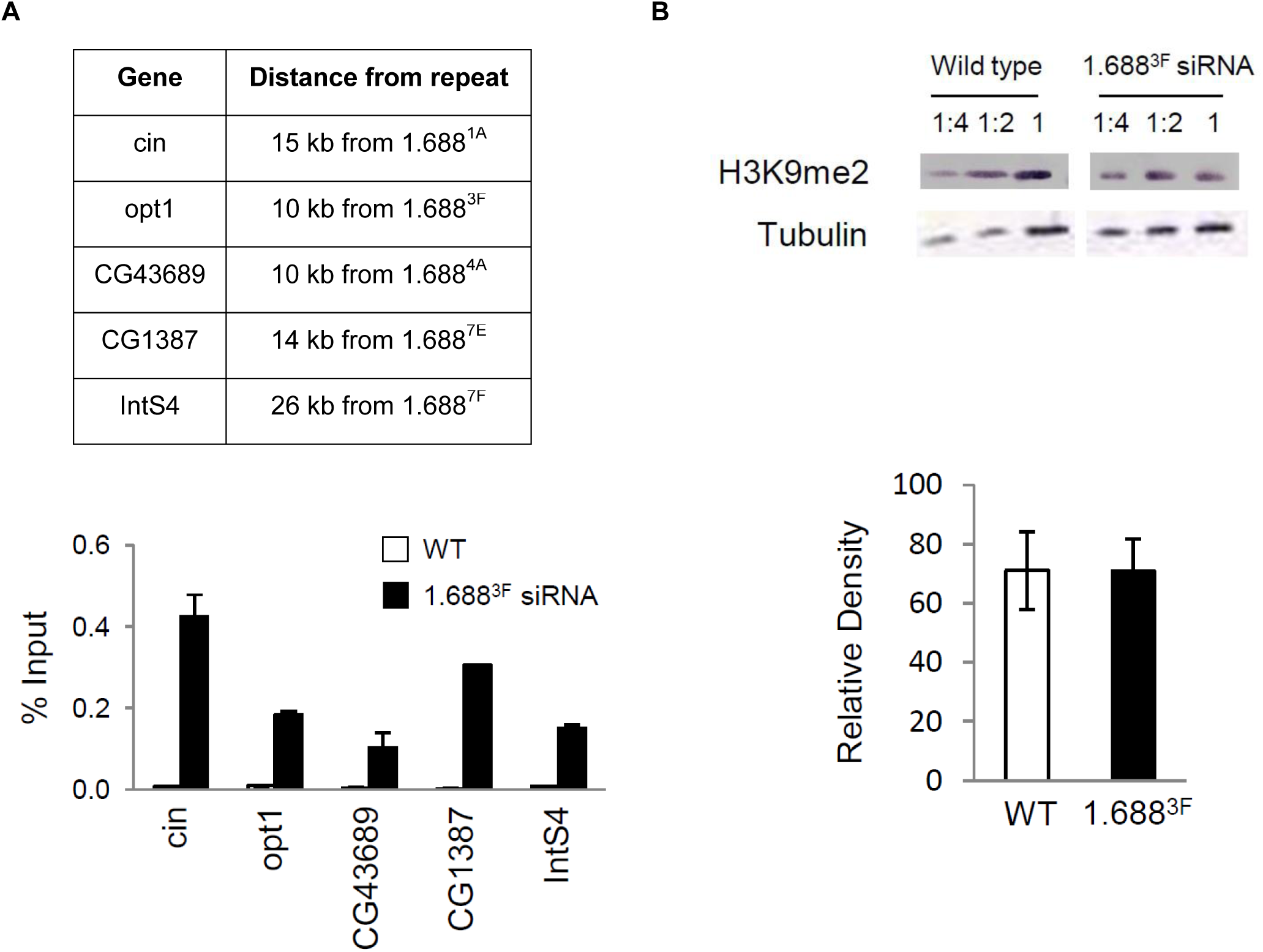
Widespread alteration in H3K9me2 around 1.688^X^ repeats is not reflected in global H3K9me2 level. (A) Genes over 20 kb from 1.688^X^ repeats display increased H3K9me2 following ectopic 1.688^3F^ siRNA production. (B) Western blot of histones from control (wild type) and 1.688^3F^ siRNA-expressing embryos does not detect a change in H3K9me2 level. H3K9me2 levels in 6-12 h embryos were compared to a tubulin loading control. Sample dilution was used to confirm signal linearity. Signal intensity was determined by ImageJ software. Standard error is derived from three biological replicates.

**Figure S4.**
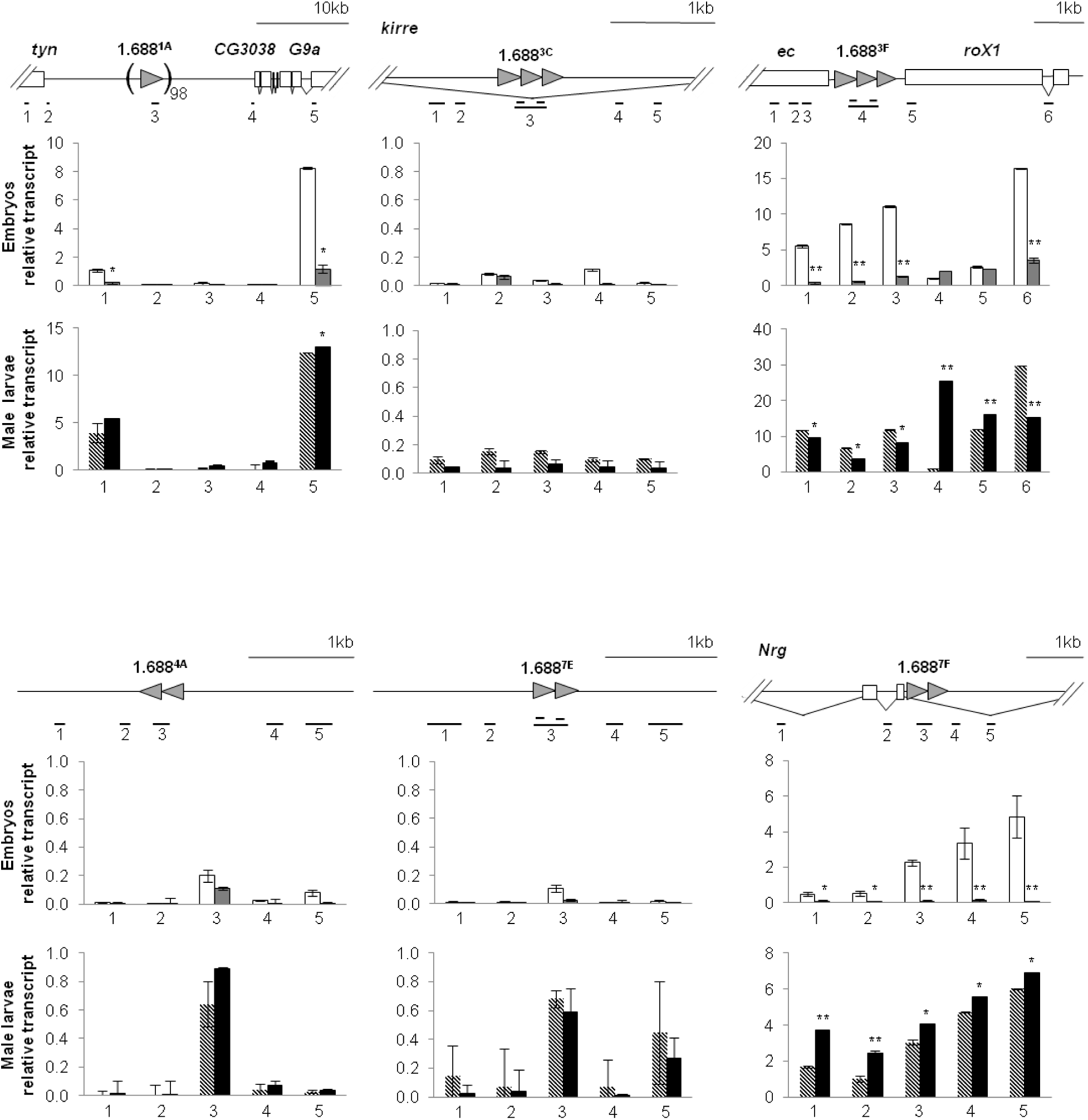
Accumulation of transcripts from 1.688^X^ repeats and surrounding regions is influenced by 1.688^3F^ siRNA. Transcript accumulation in 6-12 h embryos (white and gray bars) and male larvae (hatched and black bars) was measured by quantitative RT-PCR. White and hatched bars are controls. Gray and black bars express 1.688^3F^ siRNA. Expression is normalized to *dmn*. Standard error is derived from two biological replicates. Significance was determined using Student’s two sample *t*-test, ≤0.05 (*),≤0.001 (**) significance. Primers are presented in Table S1.

**Figure S5.**
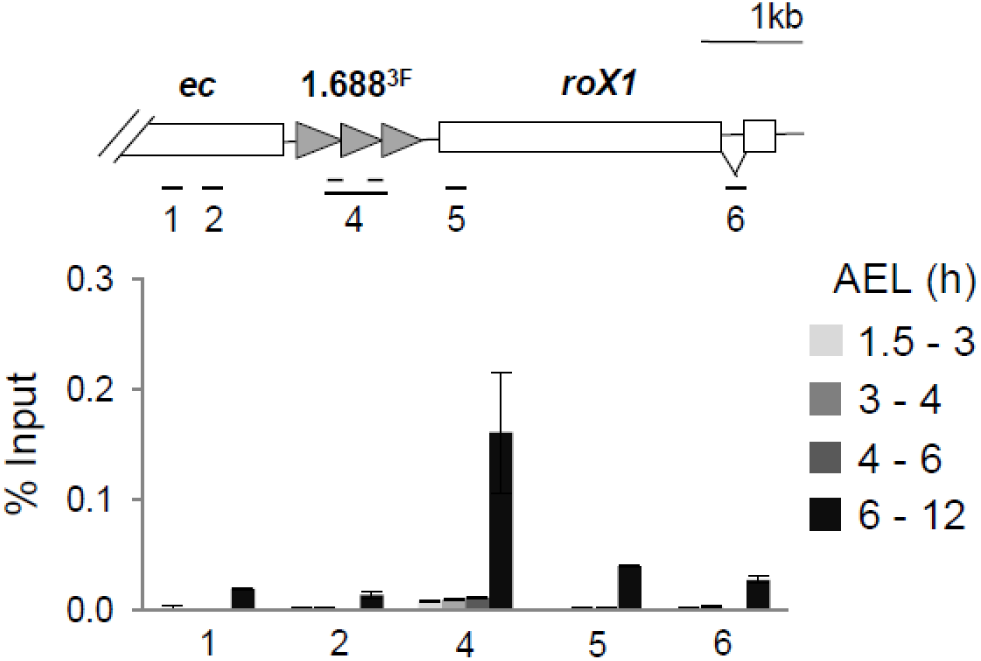
H3K9me2 deposition on 1.688^3F^ chromatin occurs 6-12 h AEL. Chromatin from staged embryos was subjected to ChIP for H3K9me2. X-localization of the MSL complex is first detected at 3 h AEL (after egg laying), but H3K9me2 enrichment is not apparent until 6-12 h AEL. Standard error is derived from two biological replicates.

**Figure S6.**
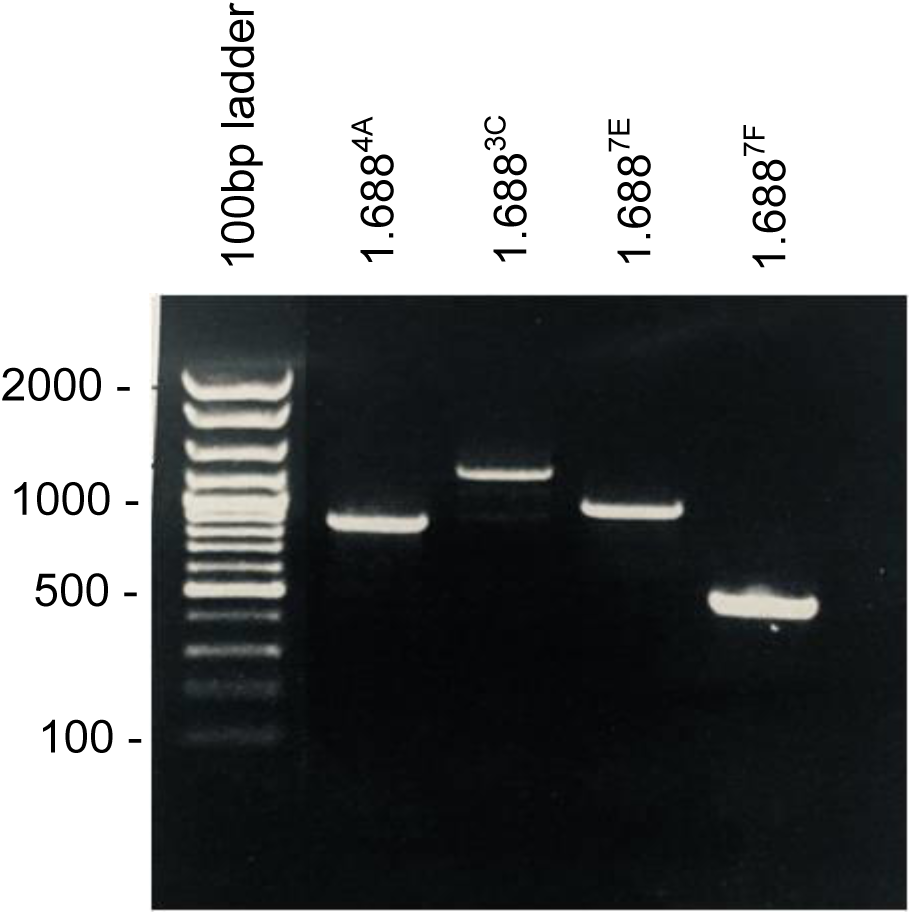
Determination of tandem repeat number in the laboratory reference strain. The size of amplicons templated with DNA from the laboratory reference *yw* strain were used to estimate the number of tandem repeats in 1.688^4A^, 1.688^3C^, 1.688^7E^, and 1.688^7F^ (see File S1).

**Table S1.**
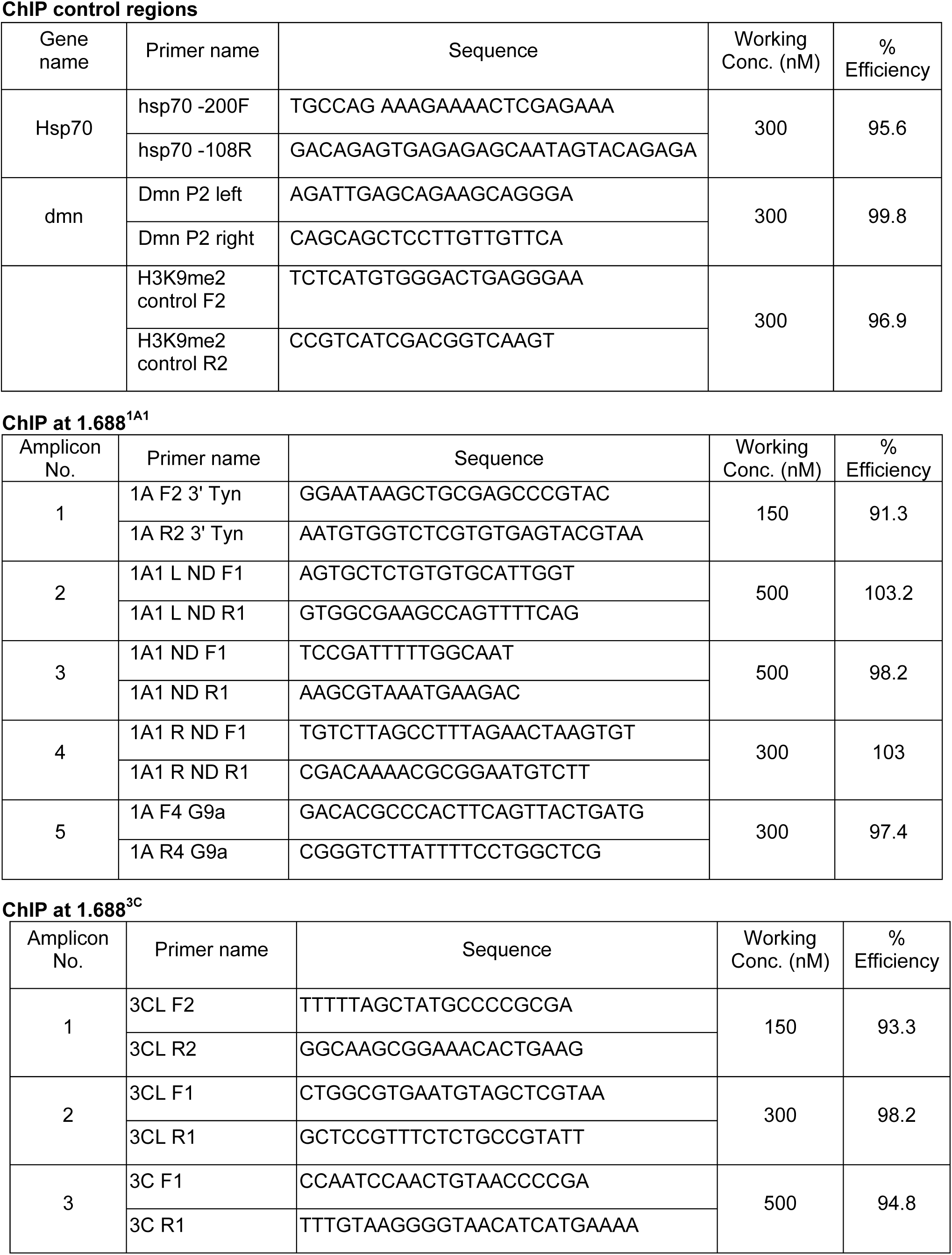
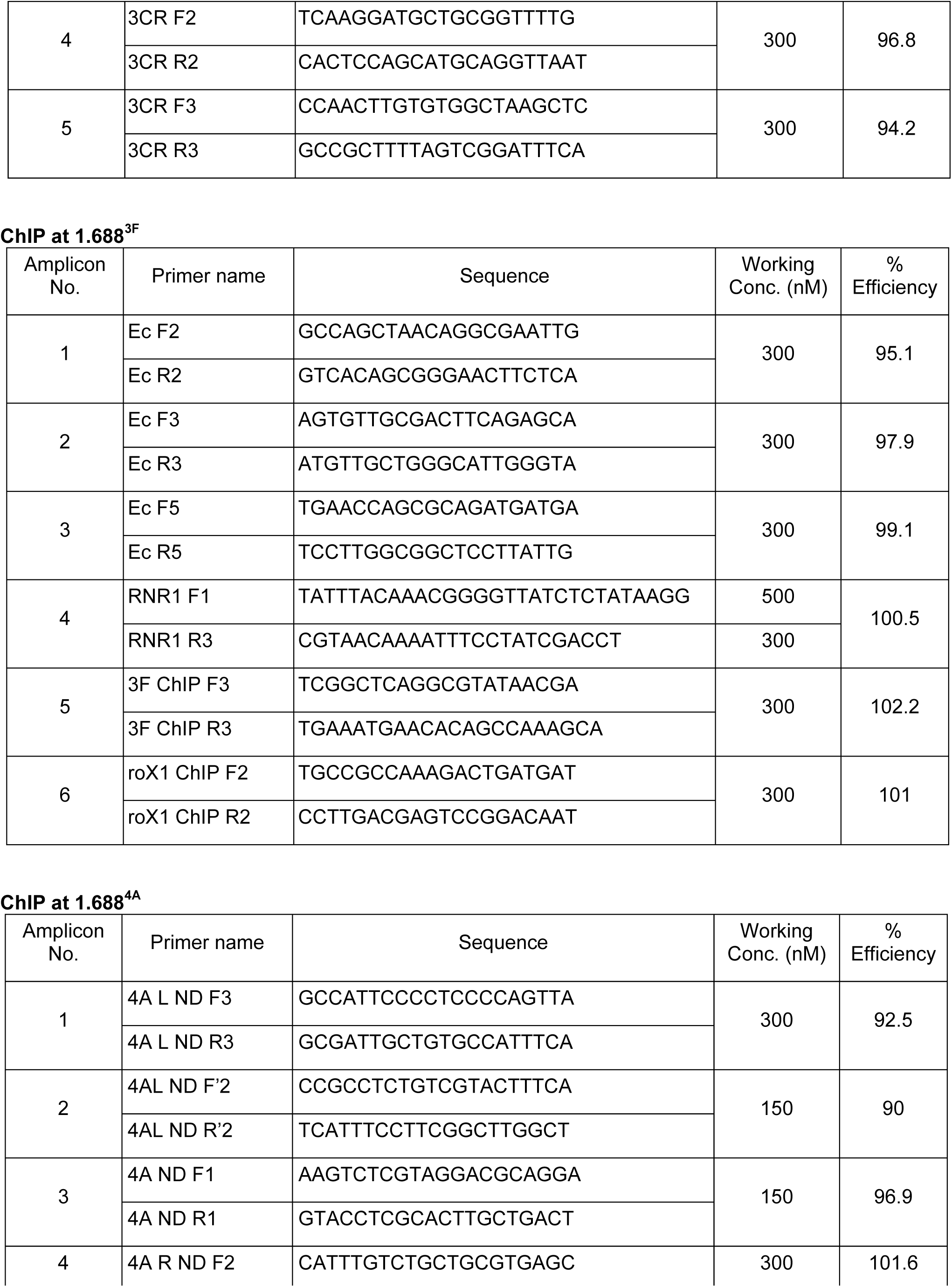
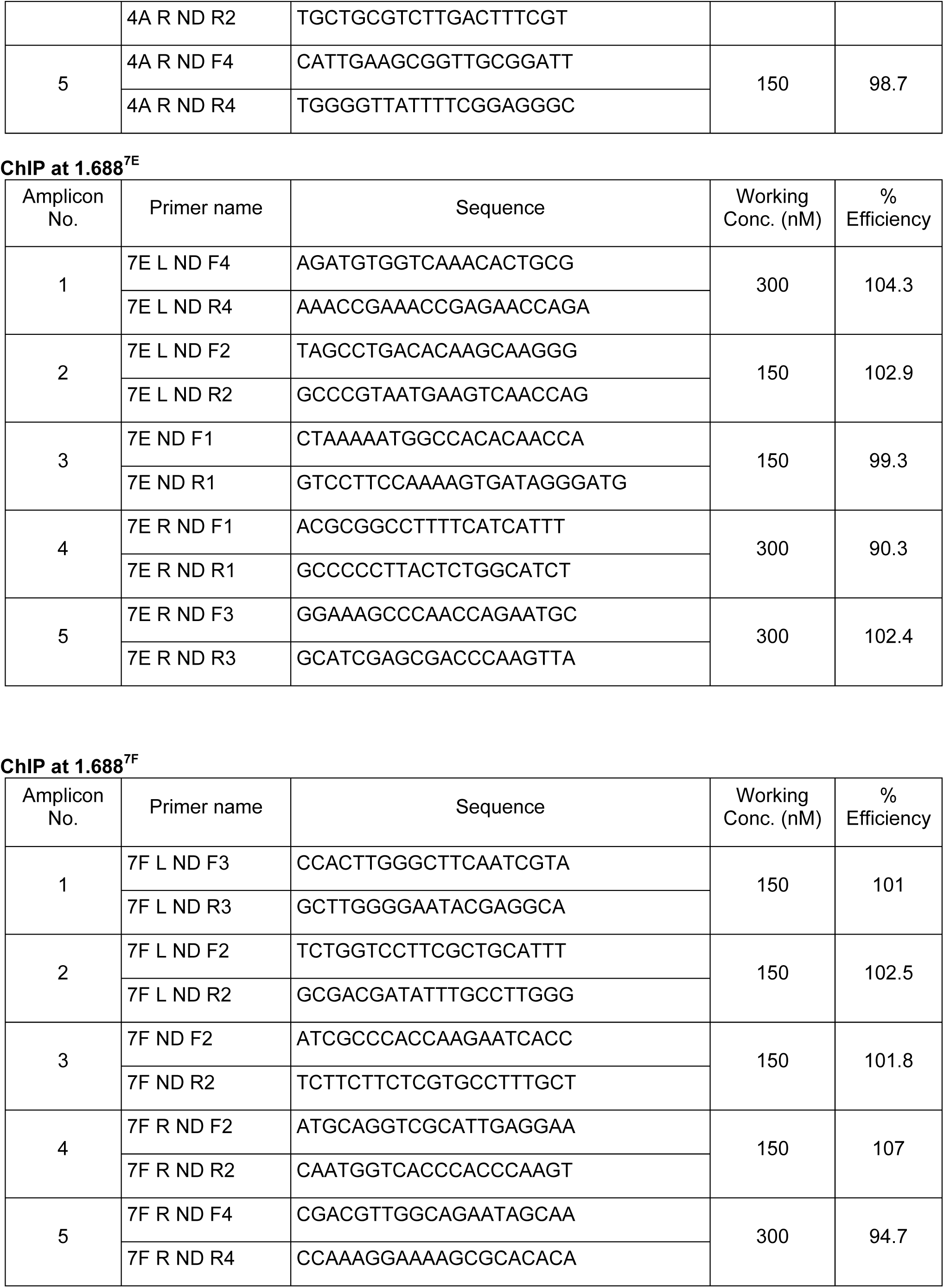
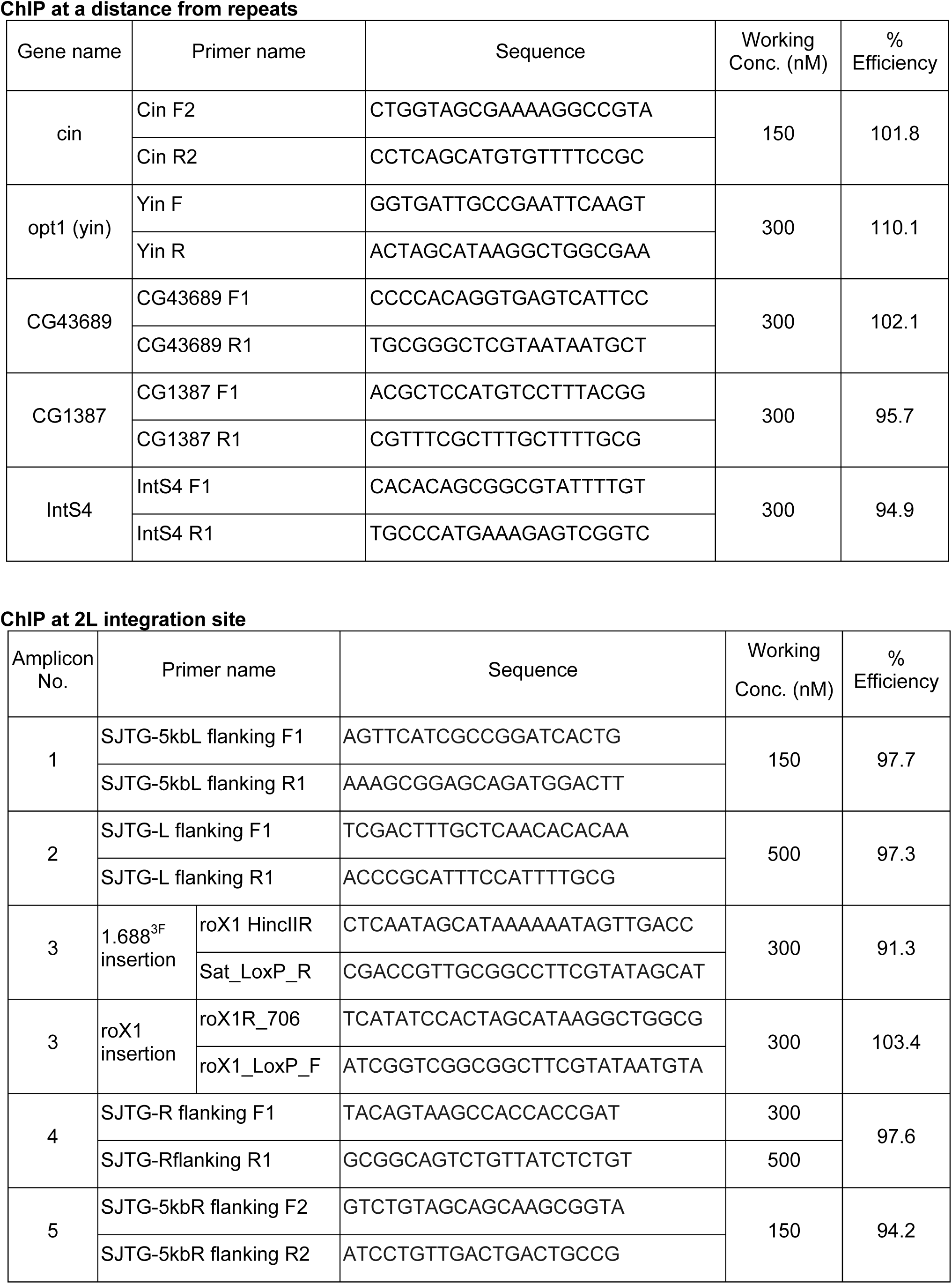
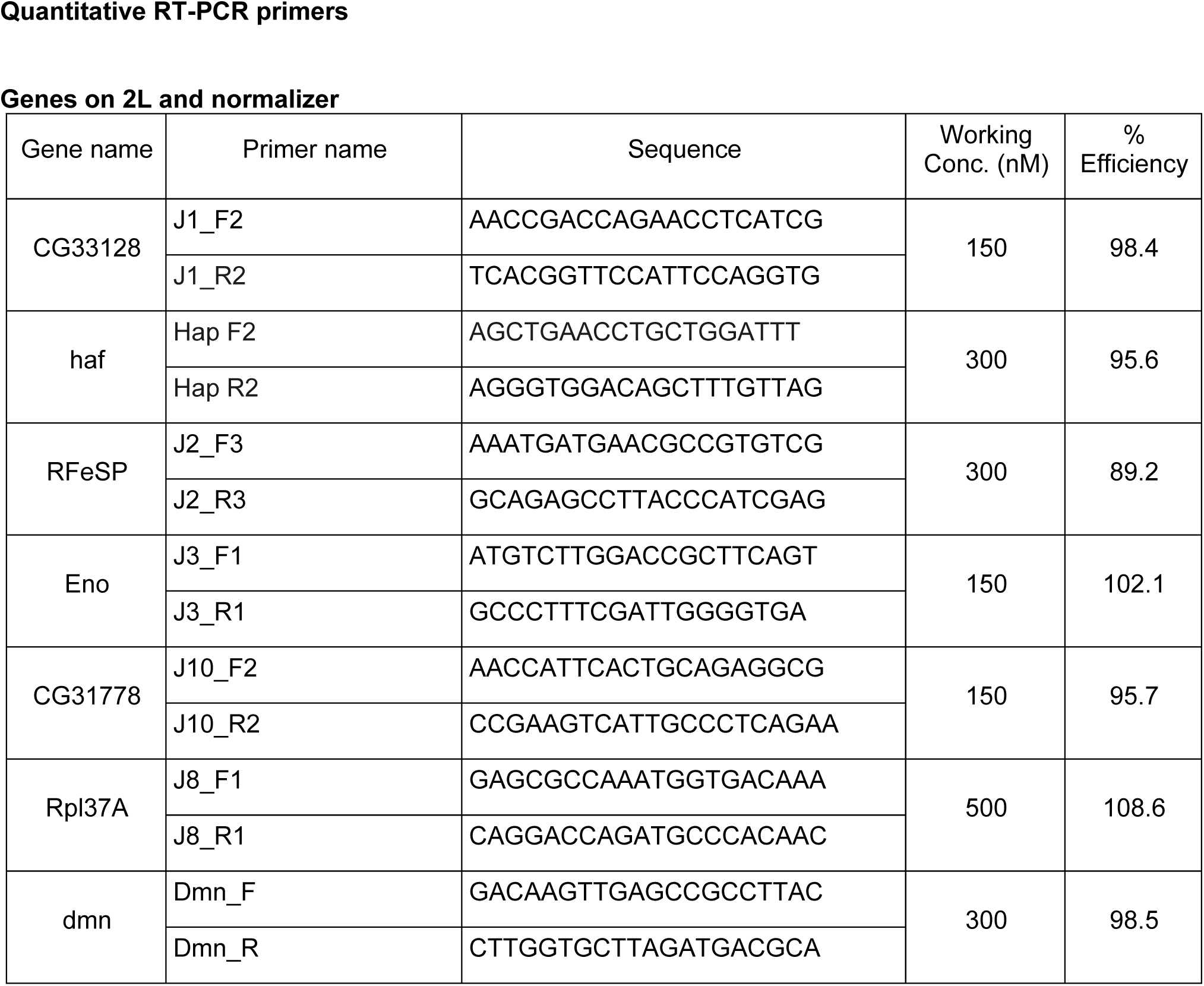
Pimers used in this study

**File S1. Estimating tandem repeat copy number in the laboratory reference strain**.

To determine the number of tandem repeats at various cytological positions in the laboratory reference (*yw*) strain, we performed PCR using primers flanking each repeat cluster. The length of the amplicon attributable to repeats was divided by 359 to obtain the copy number. Prior studies have shown that the copy number of 1.688^3F^ repeats in the laboratory reference strain is 3.5 (Menon et al., 2014). The 1.688^4A^, 1.688^3C^, 1.688^7E^, and 1.688^7F^ repeat clusters span 850 bp, 1200 bp, 900 bp, and 500 bp, respectively, bringing their copy numbers to 2.5, 3.5, 2.5, and 1.5 (Figure S6, Table 1). As the 1.688^1A^ repeats have a copy number of ∼100, it was not possible to use this method to determine 1.688^1A^ repeat length or copy number in the laboratory reference strain.

**File S2. Selection criteria for Ago2-interacting proteins**

Four criteria were used to rank Ago2-interactors in the Biological General Repository for Interaction Datasets (BioGRID) version 3.4.144, a manually curated database containing high-probability protein interactions (Stark et al., 2006), or identified by the Easy Networks (esyN) search engine (Bean et al., 2014). As BioGRID represents a subset of esyN interactions, all BioGRID interactions in Table S2 are also present in esyN. The Curation Score in Table S2 is the sum of four criteria. The method of detection was scored 0 if high throughput, and 1 if low throughput or experimentally validated. Additional criteria are: a known role in RNA interference; chromatin modification; and chromatin association. Proteins received a score of 1 for each criterion satisfied and 0 if unsatisfied. Most proteins with a score of 2 or more were tested for genetic interaction with *roX1 roX2*. Phosphatidylinositol glycan anchor biosynthesis class S (PIG-S, curation score 1) was also tested. No genes with a score ≤ 2 displayed a genetic interaction with *roX1 roX2*, but several with scores of 3 or 4 enhanced the lethality of *roX1 roX2* males, suggesting a role in dosage compensation (Table S2, Figure 2).

**Table S2.**
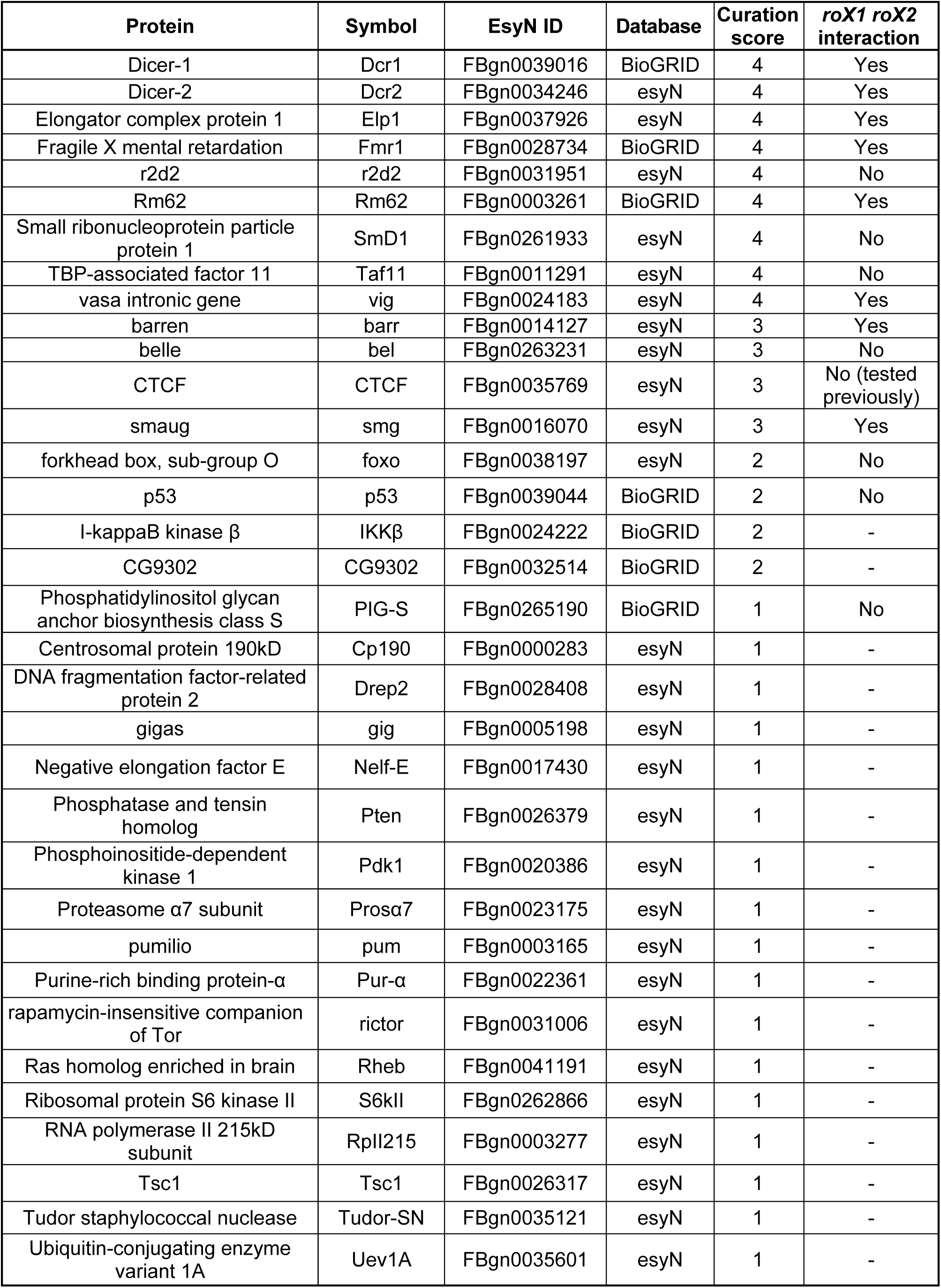
Ago2-interactors ranked by manual curation.

